# *Ret* loss-of-function decreases neural crest progenitor proliferation and restricts developmental fate potential during enteric nervous system development

**DOI:** 10.1101/2021.12.28.474390

**Authors:** Elizabeth Vincent, Sumantra Chatterjee, Gabrielle H. Cannon, Dallas Auer, Holly Ross, Aravinda Chakravarti, Loyal A. Goff

## Abstract

The receptor tyrosine kinase gene *RET* plays a critical role in the fate specification of enteric neural crest-derived cells (ENCDCs) during enteric nervous system (ENS) development. Pathogenic *RET* loss of function (LoF) alleles are associated with Hirschsprung disease (HSCR), which is marked by aganglionosis of the gastrointestinal (GI) tract. ENCDCs invade the developing GI tract, proliferate, migrate caudally, and differentiate into all of the major ENS cell types. Although the major phenotypic consequences and the underlying transcriptional changes from Ret LoF in the developing ENS have been described, its cell type and state-specific effects are unknown. Consequently, we performed single-cell RNA sequencing (scRNA-seq) on an enriched population of ENCDCs isolated from the developing GI tract of *Ret* null heterozygous and homozygous mouse embryos at embryonic day (E)12.5 and E14.5. We demonstrate four significant findings: (1) *Ret*-expressing ENCDCs are a heterogeneous population composed of ENS progenitors as well as glial and neuronal committed cells; (2) neurons committed to a predominantly inhibitory motor neuron developmental trajectory are not produced under Ret LoF, leaving behind a mostly excitatory motor neuron developmental program; (3) HSCR-associated and *Ret* gene regulatory network genes exhibit distinct expression patterns across *Ret-*expressing ENCDC with their expression impacted by *Ret* LoF; and (4) *Ret* deficiency leads to precocious differentiation and reduction in the number of proliferating ENS precursors. Our results support a model in which Ret contributes to multiple distinct cellular phenotypes associated with the proper development of the ENS, including the specification of inhibitory neuron subtypes, cell cycle dynamics of ENS progenitors, and the developmental timing of neuronal and glial commitment.

**Summary Statement:** *Ret* LoF affects proper development of the mouse ENS through multiple distinct cellular phenotypes including restriction of neuronal fate potential, disruption of ENCDC migration, and modulation of progenitor proliferation rate.

## Introduction

The enteric nervous system (ENS) is primarily derived from migrating vagal and sacral neural crest cells (Kulkarni et al. 2020; Le Douarin and Smith 1988; Kulkarni et al. 2018; Durbec et al. 1996; Kapur et al. 1992) that colonize the developing gut tube between embryonic day (E)9.5 to E14.5 in mice (Young et al. 1998) and Carnegie stage (CS) 14 to CS22 in humans (Goldstein et al. 2013), leading to the neuronal networks of the myenteric and submucosal plexuses of the gut. The ENS has essential roles in propulsive and mixing motility, absorption, immunity, and vascular function (Furness 2012) and disruption of proper ENS development or organization underlies several disorders of the gastrointestinal system (Heuckeroth 2018). Previous studies of ENS development have demonstrated that invading enteric neural crest-derived cells (ENCDCs) are predominantly neuronal precursors early in mouse development (E10.5), but as development proceeds they differentiate into either glia, expressing classical glial markers including *Gfap, Plp1,* and *S100b* (Hao and Young 2009; Kang et al. 2021), or neurons (Hao and Young 2009). Single-cell transcriptional profiling of mouse ENCDCs during mid and late (E12.5, E15.5 and E18.5) developmental time points reflects this heterogeneity and temporal change in cell fate (Lasrado et al. 2017; Morarach et al. 2021, 2017; Elmentaite et al. 2021).

One of the key factors involved in the migration of ENCDCs into the developing gut, and their differentiation into enteric neurons and glia, is the receptor tyrosine kinase RET (Natarajan et al. 2002; Lasrado et al. 2017). RET expression in ENCDCs promotes proliferation and cell survival via the MAPK and PI3K/AKT pathways (Mulligan 2014). Consequently, gain-of-function (GoF) mutations in *RET* have been associated with various cancers including medullary thyroid carcinoma and multiple endocrine neoplasias type 2A and 2B (Chatterjee et al. 2016b). Conversely, *RET* LoF due to both coding and non- coding regulatory variants leads to Hirschsprung disease (HSCR; congenital intestinal aganglionosis), an uncommon (∼1/5,000 live births) developmental disorder of the ENS characterized by absence of enteric ganglia along variable lengths of the distal bowel (Tilghman et al. 2019; Chatterjee et al. 2016a; Kapoor et al. 2015; Chatterjee et al. 2021; Alves et al. 2013). In the developing mouse gut, *Ret* deficiency leads to significant transcriptional changes in transcription factors, signaling molecules, and genes involved in specific transport and biosynthesis processes, leading to aganglionosis (Heanue and Pachnis 2006; Chatterjee et al. 2019). Tissue-level bulk transcriptomic studies in *Ret-*deficient developing mouse gut (Heanue and Pachnis 2006; Chatterjee et al. 2019)) have uncovered significant expression changes in multiple transcription factors critical to neural crest migration and ENS differentiation including *Sox10, Pax3* and *Phox2b* (Southard-Smith et al. 1998; Lang et al. 2000; Pattyn et al. 1999), as well as genes harboring mutations in HSCR patients such as *Ednrb* and *Gdnf* (Heanue and Pachnis 2006; Chatterjee et al. 2019).

Development and fate specification in the ENS occurs in conjunction with a prominent migration along the forming gut tube (Kang et al. 2021; Nagy and Goldstein 2017). As such, temporal and spatial influences affect ENCDC competence and can modulate fate decisions during development (Kang et al. 2021; Obermayr et al. 2013). While *Ret* is well known to play a significant role in the proper establishment of the ENS, the cell-type-specific effects of *Ret* LoF on normal ENS development have not yet been fully elucidated and could contribute to our understanding of normal ENS development.

To identify the cell-type-specific contributions of *Ret* to the proper organization and development of the ENS, we examined the cellular heterogeneity of the ENS during mouse gut development in the presence and absence of *Ret*. We profiled the transcriptome of 1,003 enteric neural crest-derived single cells in heterozygous *Ret* null and prenatally lethal homozygous *Ret* null mice at E12.5 and E14.5. The *Ret* heterozygous mice do not exhibit aganglionosis in the developing gut or any other known phenotypes associated with *Ret* LOF and hence serves as an suitable control for our study which focusses on the cellular dysregulation of enteric neurons and glia; the primary cell affected cell type due to *Ret* LoF. ENCDCs and their progeny were isolated using a cyan fluorescent protein (CFP) reporter driven by the *Ret* promoter for enrichment (Uesaka et al. 2008). Using a combination of differential gene expression analysis, gene co-expression pattern analysis, RNA velocity, and cell cycle state predictions, we identify several key developmental phenotypes associated with the loss of *Ret.* We find the specific effects of *Ret* LoF vary across different cellular states in ENCDCs including effects on progenitor proliferation, migration, differentiation, and fate specification. We confirm a previously described bifurcation in enteric neuronal developmental fate specification (Morarach et al. 2021; Elmentaite et al. 2021) at an even earlier time point. We further demonstrate that *Ret* expression is required for commitment to one of these two neuronal fates and subsequently, the establishment of cell types from a predominantly inhibitory neuron developmental trajectory. Additionally, we show that homozygous *Ret* LoF leads to precocious differentiation of ENCDCs and a significant reduction in ENCDC progenitor proliferation rate. Each of these observed phenotypes may independently contribute to the deficits in proper ENS cell numbers, development, organization, and function observed in HSCR. These results highlight the diverse cell-type-specific roles for *Ret*, a key developmental gene, in the proper establishment of the ENS with implications for understanding HSCR-related phenotypes, and provide insights into the unique nature of ENS development.

## Results

To assess cell-type-specific roles for *Ret* in the developing ENS, we used a previously described mouse line harboring a *Ret*-null fluorescent reporter allele (*Ret^CFP^*) (Uesaka et al. 2008). All ENCDCs express *Ret* during early development (Natarajan et al. 2002) allowing for their isolation via this reporter system. We sorted *Ret* expressing ENCDC via FACS to decipher the cell autonomous effect of *Ret* LoF. *Ret*-null (*Ret^CFP/CFP^*) mouse embryos exhibit total colonic aganglionosis and kidney agenesis, and are perinatal lethal (Uesaka et al. 2008). Importantly, no defects in ENS formation have been observed in *Ret*-heterozygous (*Ret^CFP/+^)* mice (Uesaka et al. 2008) nor any significant difference in the genome wide expression observed when compared to wildtype GI tract at the same developmental times (Kapoor et al. 2017). Comparison of *Ret^CFP/CFP^* to *Ret^CFP/+^* mice can therefore elucidate the mechanisms by which *Ret* LoF affects the proper formation of the ENS.

To investigate transcriptional changes of ENCDCs, we performed single-cell RNA-sequencing (scRNA-seq) on FACS-enriched populations of ENCDCs in the developing murine gut. Briefly, CFP- positive single cells were collected from the dissociated stomach and intestine of male and female E12.5 and E14.5 *Ret^CFP/+^* and *Ret^CFP/CFP^* embryos and subjected to a modified Smart-seq2 scRNA-Seq library preparation (Picelli et al. 2014; Chevée et al. 2018) (see Methods). Libraries from 95 cells for each of two biological replicates were sequenced from all age, sex, and genotype combinations, with the exception of E14.5 *Ret^CFP/CFP^* embryos. Consistent with a previously described reduction in the number of CFP^+^ cells seen in a *Ret* hypomorphic mouse model (Uesaka et al. 2008), we observed reduction in the cell capture rate in the E14.5 *Ret^CFP/CFP^* mice as compared to other ages and genotypes. We also observed a pronounced sex bias in the number of CFP^+^ cells recovered from E14.5 *Ret^CFP/CFP^* embryos, with fewer viable cells collected from males, consistent with the established sex bias observed in HSCR (Amiel et al. 2008) and described in other mouse models of HSCR (Dang et al. 2011; Uesaka et al. 2008). This reduced number of Ret^CFP/CFP^ cells as a function of developmental age and sex, highlights a specific challenge of this mouse model as the number of viable CFP^+^ cells rapidly diminishes in these conditions. As such, only 22 cells from one male E14.5 *Ret^CFP/CFP^* embryo and 190 cells from four female E14.5 *Ret^CFP/CFP^* embryos were recovered (Table 1). Full-length cDNA scRNA-Seq libraries from 1,341 individual cells were sequenced to an average depth of 9.91x10^5^ (standard deviation 7.43x10^5^) paired- end reads per cell. Low quality cells were removed using standard metrics (see Methods) leaving 1,003 cells meeting all quality control parameters (Figure 1A; Table 1) with an average read depth of 1.16x10^6^ (standard deviation 6.83x10^5^) paired-end reads per cell. Relative FPKM values were converted to estimated RNA copies per cell (CPC) using the CENSUS algorithm (Qiu et al. 2017). We identified a set of 14,319 expressed genes with greater than zero CPC in at least 20 cells and a standard deviation greater than zero across all cells. To visualize the transcriptional relationships between cells, we identified a set of 419 high-variance genes to use as input to the uniform manifold approximation and projection (UMAP) dimensionality reduction tool to establish a biologically relevant two-dimensional embedding of the cells (Figure 1B, Figure S1).

**Figure 1:**
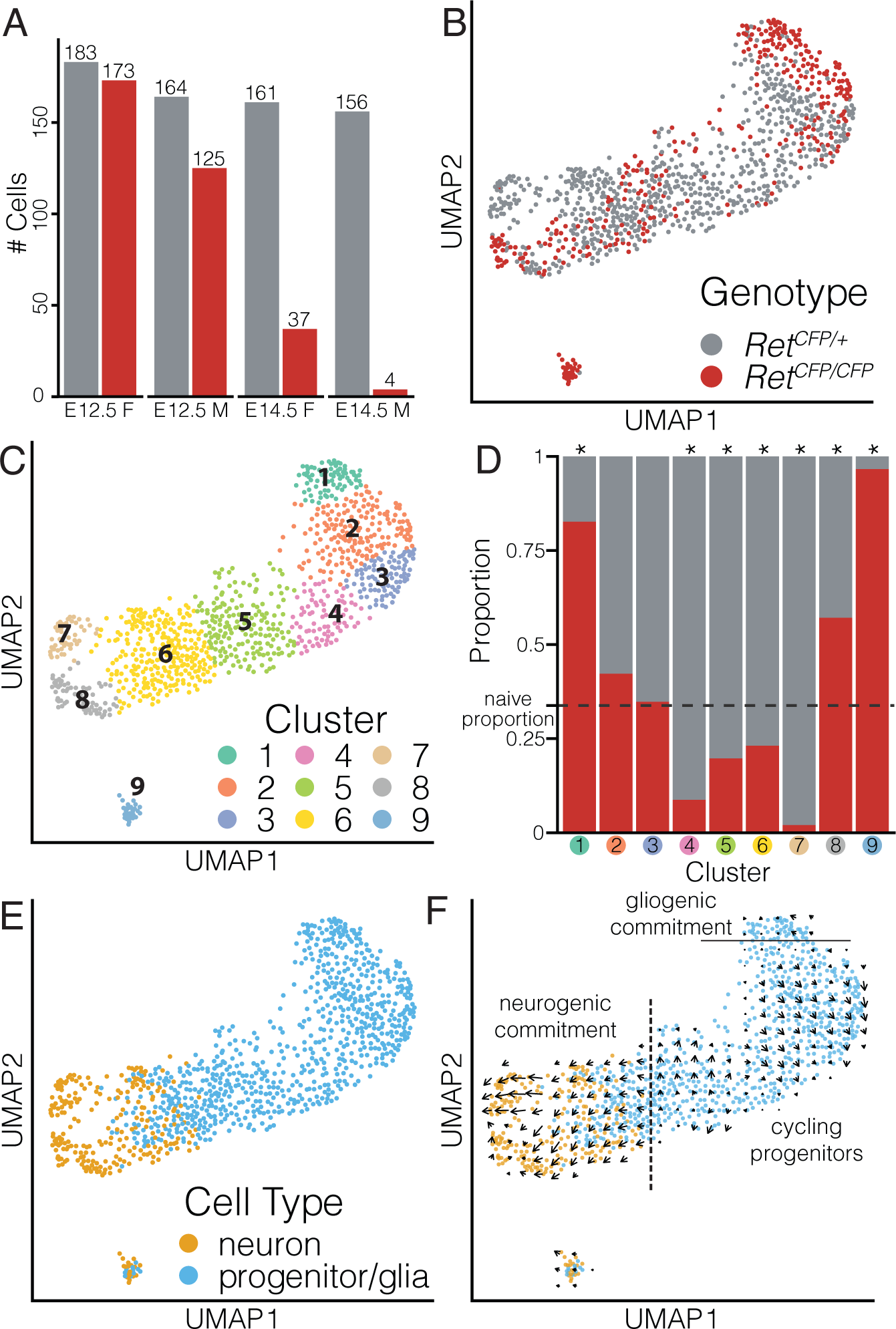
A cellular map of Ret expressing enteric nervous system cells. (A) Barplot of post quality control (QC) count of sorted *Ret^CFP/+^* and *Ret^CFP/CFP^* expressing cells from E12.5 & E14.5 embryonic gut in male and female mice. (B) Uniform Manifold Approximation and Projection (UMAP) of 1003 cells colored by genotype. (C) UMAP embedding colored by 9 learned clusters (D) Stacked barplot of the genotype proportions for each cluster. Asterisk denotes significant (Chi-squared test for equal proportions, p<0.05 after multiple testing correction) deviation from expected naive proportion. (E) UMAP embedding of *Ret*-expressing cells classified as glia/progenitor or neuron (F) RNA velocity vector field along the UMAP embedding demonstrates direction of cellular differentiation. Dotted line indicates neurogenic commitment, solid line indicates gliogenic commitment.

### Annotation of cell types and states of the developing ENS

To quantify cell type and cell state differences with respect to genotype, we first used a Gaussian- based clustering algorithm (Scrucca et al. 2016) to identify 9 clusters (Figure 1C) and calculated gene- wise specificity scores to identify gene expression profiles for each cluster (Figure S2A). High specificity scores relative to mean expression identified clusters of actively cycling progenitors (cluster 2, Hist1h family genes; cluster 3, *Aurka/b*, *Cdk1*), a neurogenic commitment window (cluster 4, *Neurod4*; cluster 5, *Pcp2, Hes5*, cluster 6, *Hes6*, *Btg2*), maturing neurons (cluster 7, *Etv1*, *Vip*, *Gal, Cartpt*; cluster 8, *Slc18a3 (ChAT), Npy, Snap25, Gfra2*; cluster 9, *Th, Dbh, Ddc*), as well as a population of maturing enteric glia (cluster 1, *Plp1*, *Myl9*) (Figure S2B, Table 2). The neural progenitor cell markers p75 (*Ngfr*) and the pan neuronal marker TuJ1 (*Tubb3*), which have traditionally been used to label mature enteric neurons along with *Ret* (Hao and Young 2009; Young et al. 1999), have a distinct, largely non-overlapping expression profile in *Ret-*expressing cells at these time points (Figure S2C). Consistent with this, we observe *Ngfr* and *Tubb3* expression to be restricted to non-overlapping populations of cells at either end of our learned manifold.

To annotate individual cells as either progenitors, developing glia, or developing neurons we next evaluated the use of canonical marker gene expression for mature ENS cell types. However, during development, the majority of cells may not yet express markers of mature, terminally differentiated cell types. To address this limitation, we chose to assign cell types using a combination of transfer learning and marker gene expression, where available, to predict the most likely annotation probabilities based on an existing cellular atlas of adult mouse enteric neurons and glia (Zeisel et al. 2018). To obtain transcriptional signatures of mature ENS cell types, we learned patterns of gene co-expression on the annotated set of enteric neurons and glia from the public dataset using single-cell Coordinated Gene Activity in Pattern Sets (scCoGAPS) (Stein-O’Brien et al. 2019) and trained a random forest (RF) model on those learned patterns to predict cell type (see Methods). We next projected our single-cell data into the latent spaces defined by these patterns using projectR (Stein-O’Brien et al. 2019; Sharma et al. 2019) (Figure S3) and employed the RF model to classify cells in our dataset based on their projected pattern weights, thereby agnostically annotating the cells in our data as glia or neuron (Figure S4A-C). We next calculated cell type proportions by cluster (Figure S4D) and for clusters in which at least 90% of cells shared an annotation, we transferred this majority annotation to all members of a given cluster. As a result, clusters 1-5 (1 = 99% glia, 2 = 98%, 3 = 92%, 4 = 95%, 5 = 94%) were annotated as glia and clusters 7 and 8 (7 = 98% neuron, 8 = 90%) were annotated as neurons. Annotations for clusters 6 (58% glia) and 9 (63% neuron) were not altered.

Our single-cell gene expression profiling is a cross-sectional snapshot in developmental progression for a population of cells that have not yet fully committed to mature cell type identities. To infer the direction of state transitions for our ENCDC population, we next calculated RNA velocity estimates (La Manno et al. 2018) and observed that the learned velocity fields comprise diverging trajectories from a common origin (Figure 1F). Cells with high projection weights in patterns associated with proliferating ENCDCs (pattern 40: *Brca1/2, Cdc6, Pola1/2*; pattern 41: *Ccna1/b1, Cenpe, Kif24*; pattern 42: *Aurka/b, Bub1, Top2a*; Figure S3, Tables 3, 4, and 5) exhibit an ellipsoid pattern of velocities, consistent with transition through the cell cycle. Extending from this progenitor population we observed an population of cells expressing canonical markers consistent with progression towards a neuronal fate (cluster 6: *Btg2*, *Hes6* Figure S2B) that further branches into two distinct trajectories of neuronal specification (Figure 1F), consistent with previous observations of postmitotic branching of cellular fate specification for neural crest-derived neurons in later gut development (Morarach et al. 2021). We infer through shared marker gene expression that cluster 7 in our data corresponds to a previously described developmental branch (branch A) (*Etv1^+^*, *Nos1^+^*, *Vip^+^*, *Gal^+^*) and cluster 8 corresponds to branch B (*Bnc2^+^*, *Dlx5^+^*, *Mgat4c^+^*, *Ndufa4l2^+^*) (Figure S6A). These data highlight that *Ret*-expressing cells isolated from the GI tract encompass two broad fates during development (glial-commitment and neuronal-commitment) and confirm that neuronally-committed ENCDCs diverge post-mitotically into two distinct developmental trajectories, outlining the diversity of cellular fate trajectories accessible to ENCDCs within the developing mouse gut.

### Ret LoF induces distinct transcriptional responses in different ENS cell states

We first asked whether loss of *Ret* changes the proportion of ENCDC subtypes in our dataset. To assess this, we tested for genotype bias in cluster composition by comparing the proportion of Ret^CFP/CFP^ cells against the naïve (overall) proportion of 34%. Using a chi-squared test for equal proportions, we observed significant deviations in seven of the nine identified clusters after Bonferroni correction for multiple tests: clusters 1, 8, and 9 were enriched for *Ret^CFP/CFP^* cells (cluster 1: 83%, Bonferroni adjusted p=9.86x10^-18^; cluster 8: 57%, p=5.64x10^-4^; cluster 9: 97%, p=1.24x10^-11^), clusters 4-7 were depleted for *Ret^CFP/CFP^* cells (cluster 4: 9%, Bonferroni adjusted p=3.48x10^-5^; cluster 5: 20%, p=1.00x10^-3^; cluster 6: 33%, p=3.44x10^-3^; cluster 7: 2%, p=6.33x10^-5^), while clusters 2 and 3 were relatively invariant (cluster 2: 42% *Ret^CFP/CFP^*, Bonferroni adjusted p=0.196; cluster 3: 35%, p=1.0) (Figure 1D). Clusters 6, 7, and 8 are neuronal-committed clusters and differ significantly from the naive proportion. Clusters 6 and 7 are enriched for *Ret^CFP/+^* cells and 8 is enriched for *Ret^CFP/CFP^* cells, indicating strong genotypic effects associated with neuronal fate specification. Three glial clusters, clusters 1, 4, and 5, differ significantly from the naive proportion as well, with cluster 1 enriched and clusters 4 and 5 depleted for *Ret^CFP/CFP^* cells. Cluster-specific enrichment of genotype within the neuronal and glial populations suggests that genotypic effects of *Ret* LoF modulates cell fate commitment with divergent results across ENCDC cell states.

To identify the transcriptional effects of *Ret* LoF on the developing ENS, we tested 14,319 genes and identified 520 as differentially expressed with respect to genotype across all cells (monocle likelihood ratio test; Benjamini-Hochberg corrected q-value < 0.05, Figure 2A, Table 6A). Hierarchical clustering based on expression of these 520 genes revealed that cells cluster first by cell type then by genotype, again suggesting that the transcriptional changes observed are largely cell-type-specific. Gene ontology analysis revealed enrichment for several relevant processes, including gliogenesis, cell fate commitment, and axon development (hypergeometric test q-value < 0.05, Figure 2B, Table 7A). Significantly upregulated genes expressed in *Ret^CFP/CFP^* progenitor/glia includes several associated with ENS glial maturation (*Fabp7*, *Plp1*, *Myl9*), indicating that these progenitor/glia cells may be on average more mature than their *Ret^CFP/+^* counterparts. A second cluster of genes is more highly expressed in *Ret^CFP/+^* neurons and includes genes with specific expression in the neurogenic branch A (*Cartpt, Vip, Nos1, Etv1*), consistent with our observation that *Ret*^CFP/+^ cells are significantly overrepresented in branch A (cluster 7).

**Figure 2:**
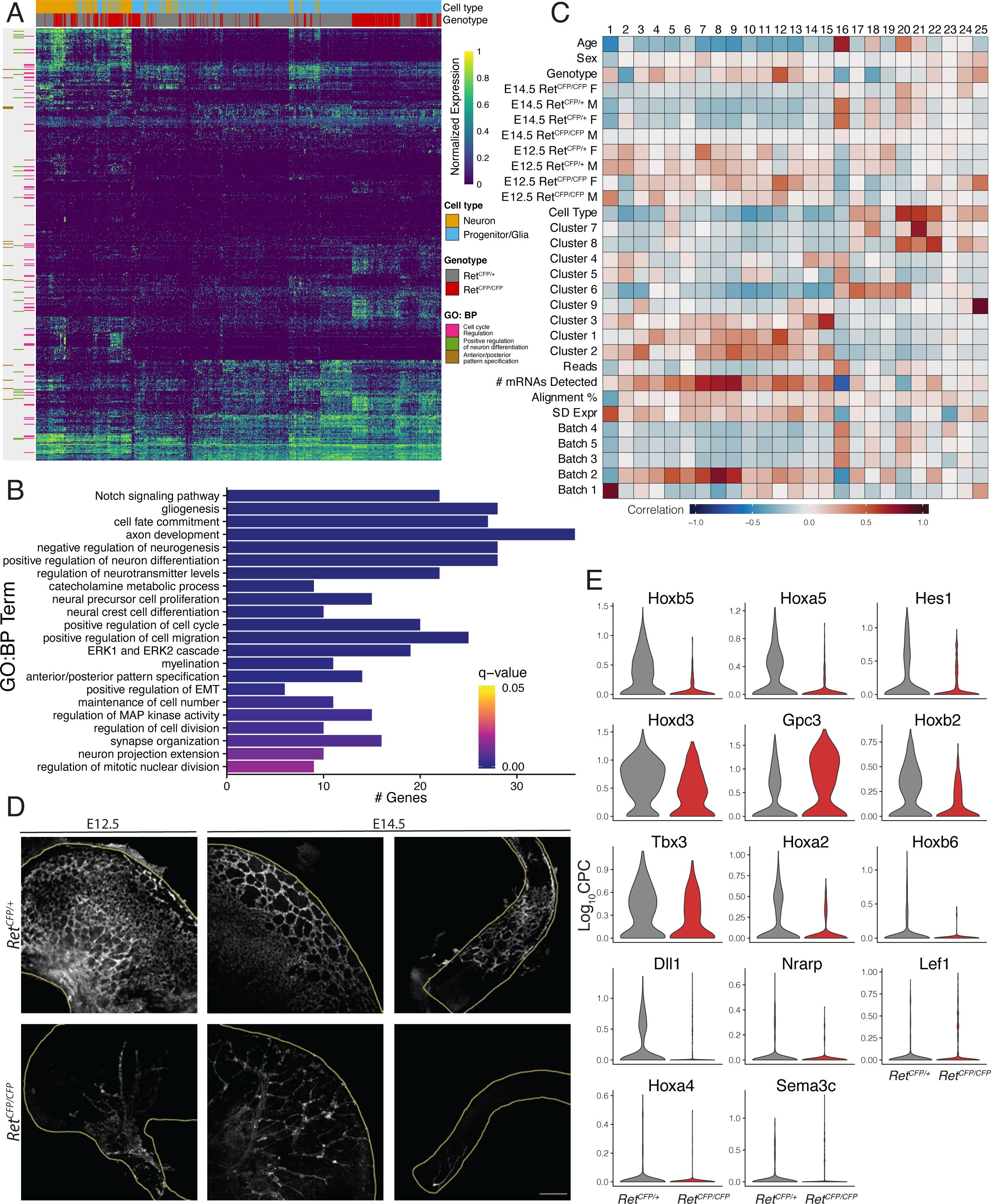
Cellular and molecular changes driven by Ret loss of function. (A) Heatmap of differentially expressed genes between *Ret^CFP/+^* and *Ret^CFP/CFP^* cells. (B) Significant (hypergeometric test, p<0.05) gene ontology (GO) gene sets include neuron differentiation, gliogenesis and cell fate commitment. C) Correlation plot of scCoGAPS patterns vs. annotated cellular features including developmental age, sex and genotype. (D) Immunofluorescence staining of mouse GI tract at E12.5 and E14.5 showing the organization of CFP-expressing cells. The first two columns show stomach and the third column shows hindgut. 10X magnification, scale bar is 200 μm. (E) Violin plots of significantly differentially expressed genes associated with the anterior-posterior patterning GO term.

We next sought to map the transcriptional signatures learned from our ENCDC dataset to specific cellular features by using coordinated sets of co-regulated genes. To this end, we applied scCoGAPS to learn 25 patterns of gene co-expression from our ENCDC scRNA-Seq dataset (Figure S5, Table 8). We calculated the Pearson correlation for pattern weights by cell against both known and learned cellular annotations (Figure 2C, Table 9). This revealed patterns that correlated with technical (read depth and alignment rates), experimental (genotype, age, and sex), as well as biological (cell type and cluster) variables. We identified patterns correlated with the acquisition of different neuronal identities (Pattern 20: *Elavl4* (*HuD*)*, Nrxn3, Tubb3, Nnat, Syt1*; Pattern 21: *Nos1, Tubb2a, Mapt, Sncg, Ntng1*; Pattern 22: *Onecut2, Gabrg2, Gabra3, Slc18a3* (*ChAT*)*, Gfra2, Elavl2* (*HuB*)*, Elavl3* (*HuC*)) (Figure S5; Table 10), a transient neurogenic window (Pattern 19: *Pou4f1, Gadd45g, Neurod4, Hes6, Btg1/2, Sox11, Phox2b*), as well as glial identity (Pattern 12: *S100b, Plp1, Fabp7*). In addition, we also identified patterns well correlated with specific genotype-biased clusters of cells including Pattern 18 (*Ret, Gal, Ass1, Etv1*) and Pattern 21 specifically used by Branch A neurons in cluster 7, and Pattern 12, specifically used by cells in cluster 1 undergoing glial maturation and significantly enriched for *Ret^CFP/CFP^* cells.

Previous in vivo studies in mice have shown a severe migratory delay beginning as early as E9.5 in the absence of *Ret (Natarajan et al. 2002; Uesaka et al. 2008, 2015)*. Consistent with this observation, we identified four gene expression patterns (Patterns 2, 18, 19, and 25), well correlated with genotype, where *Hox* gene family members were identified as pattern markers (Stein-O’Brien et al. 2017) (Table 10). Pattern 25 markers include the more distal *Hoxa/c9* and *Hoxa/c10*, and the separation of these cells from the main manifold of ENCDCs, combined with the specific expression of marker genes such as *Maob* and *Pcdh10*, suggests that the handful of cells specifically using Pattern 25 are likely caudal Schwann cell precursors. The enrichment for Hox genes within the remaining three pattern marker gene lists with strong correlation to genotype, suggests that Hox gene expression in ENCDCs may be a discriminating feature between the two genotypes. This was confirmed by the identification of seven *Hox* genes, (*Hoxa2*, *b2*, *d3*, and the more caudally-expressed *Hoxa4*, *a5*, *b5*, and *b6*), as well as several anterior-posterior pattern specification genes (*Dll1, Gpc3, Hes1, Lef1, Nrarp, Sema3c,* and *Tbx3*), significantly differentially expressed (monocle likelihood ratio test; q<0.05) with respect to genotype (Figure 2A-B). With the exception of *Gpc3, Lef1,* and *Sema3c*, all of the differentially expressed genes associated with anterior-posterior positioning show increased expression in *Ret^CFP/+^* cells (Figure 2E). Consistent with this and other previous work at E12.5 (Hirst et al. 2018), we observe that ENCDCs in the absence of *Ret* have not left the extrinsically innervating nerve fibers of the stomach by E12.5 nor by E14.5 (Figure 2D), in stark contrast to *Ret^CFP/+^* cells which have formed a dense network throughout the stomach and all but the end of the hindgut by E12.5. As the *Hox* gene family is well-known to contribute to axial patterning, this differential expression of *Hox* genes may therefore be a consequence of significant rostro-caudal positional differences in ENCDCs in the presence and absence of *Ret*.

### Ret is required for commitment to the branch A neuronal developmental trajectory

Recent work in E15.5 mouse gut shows that after neuronal commitment, cells of the developing ENS branch into two main trajectories, referred here as branch A (*Etv1*+*, Gal+, Nos1*+, *Vip*+) and branch B (*Bnc2*+, *Dlx5*+, *Mgat4c*+, *Ndufa4l2*+) (Morarach et al. 2021). Our UMAP embedding reveals this branching behavior can be observed as early as E12.5 and that the choice of branch is *Ret*-dependent (Figure 3A, Figure S6A). As observed above, cluster 7, corresponding to branch A, is almost exclusively composed of *Ret^CFP/+^* cells. Conversely, while cluster 8 (branch B) is significantly enriched for *Ret^CFP/CFP^* cells (Figure 1D), both genotypes are represented along the branch B trajectory. This suggests that there is a significant bias in the choice of developmental trajectory mediated in part by *Ret* expression in ENCDCs.

**Figure 3:**
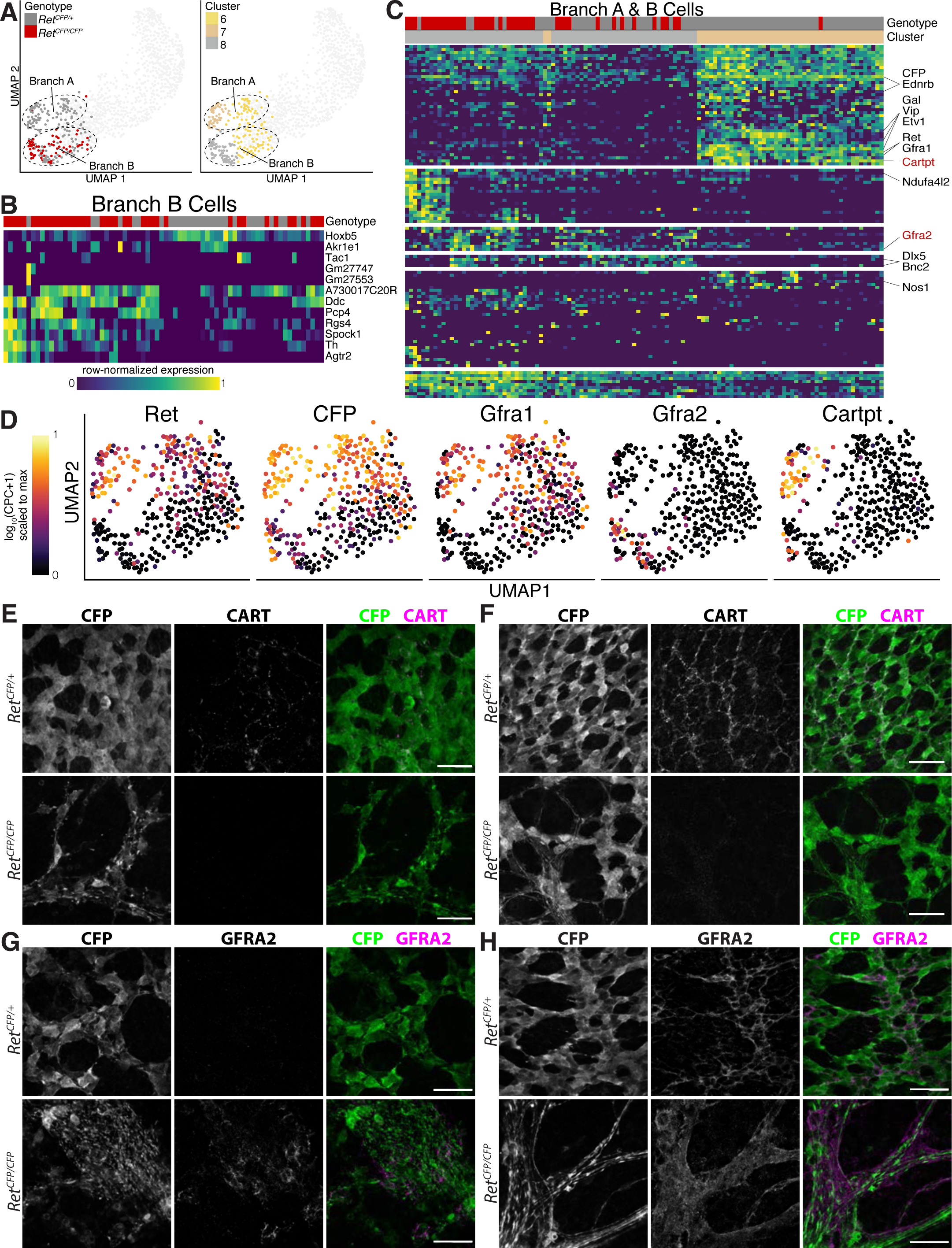
RET expression is required for access to the Branch A developmental trajectory. (A) UMAP embedding of the neuronal cells comprising both developmental branches colored by genotype (top) or by cluster assignment (bottom). Branch A (cluster 7) is almost entirely *Ret^CFP/+^* cells while both genotypes are represented in branch B (cluster 8). (B) Heatmap of significant differentially expressed genes with respect to genotype for cells in branch B. (C) Heatmap of significant differentially expressed genes with respect to developmental lineage (branch) across all neurons from clusters 7 and 8. Row cluster segmentation represents the top six divisions of the hierarchical clustering of gene expression values. (D) UMAP embeddings highlighting the persistent expression of *Ret, CFP,* and *Gfra1* along the branch A developmental trajectory, the specific induction of the non-canonical Ret co-receptor *Gfra2* along cells in branch B, and the specific expression of the branch A marker gene *Cartpt*. (E,F) Immunofluorescence assay of branch A marker CART at E12.5 in *Ret^CFP/+^* and *Ret^CFP/CFP^* stomach (E) and E14.5 in *Ret^CFP/+^* hindgut and *Ret^CFP/CFP^* stomach (F) demonstrates presence of CART positive cells only in the *Ret^CFP/+^* mice. (G,H) Immunofluorescence assay of branch B marker GFRA2 at E12.5 in *Ret^CFP/+^* and *Ret^CFP/CFP^* stomach (G) and E14.5 in *Ret^CFP/+^* hindgut and *Ret^CFP/CFP^* stomach (H). All IF assay images are 60X magnification and scale bars are 40 μm.

We next sought to identify branch-specific differences in gene expression with respect to genotype. When we tested cluster 8 (branch B), we identified 12 genes differentially expressed with respect to genotype (monocle likelihood ratio test, q<0.05, Figure 3B, Table 6B), four of which are more highly expressed in *Ret^CFP/+^* and eight more highly expressed in *Ret^CFP/CFP^*. Gene ontology analysis indicates that genes upregulated in *Ret^CFP/CFP^* are enriched (hypergeometric test, q<0.05, Table 7B) for dopamine biosynthetic processes (q<4.52x10^-6^, *Agtr2*, *Ddc*, *Th*), catecholamine biosynthetic processes (q<6.44x10^-6^, *Agtr2*, *Ddc*, *Th*), and regulation of corticosteroid hormone secretion (q<3.35x10^-4^, *Agtr2*, *Tac1*). This suggests that, while both genotypes exhibit generally similar transcriptional profiles within branch B, there may still be functional consequences of *Ret* LoF within this trajectory.

To examine differences between branch A and B trajectories and identify branch-specific markers, we tested for differential expression irrespective of genotype to identify 111 genes (monocle likelihood ratio test, q<0.05, Figure 3C, Table 6C). Gene ontology enrichment analysis (Table 7C) reveals that genes with higher expression in cluster 7 are enriched for regulation of neurological system processes (hypergeometric test, q=1.91x10^-5^, *Cartpt, Ednrb, Fgfr1, Nos1, Nrg1, Vip*) and axonogenesis (hypergeometric test, q=6.11x10^-5^, *Ablim1, Auts2, Cdh11, Etv1, Lrrc4c, Lrrn3, Nrg1, Ntng1, Ret*), while genes more highly expressed in cluster 8 are enriched for catecholamine biosynthetic processes (hypergeometric test, q=1.15x10^-8^, *Agtr2, Dbh, Ddc, Gata3, Gch1, Th*), regulation of neurotransmitter levels (hypergeometric test, q=2.00x10^-6^*, Chrna7, Dbh*, *Ddc, Gch1, Nrxn1, Prima1, Slc18a1, Sytl4, Th*), and dopamine biosynthetic processes (hypergeometric test, q=2.36x10^-6^*, Agtr2, Dbh*, *Ddc, Gch1, Th*). The increased expression of genes in this latter geneset in *Ret*^CFP/CFP^ cells in cluster 8 relative to *Ret*^CFP/+^ cells (Figure 3B) suggests that differentiation and/or maturation rate of cells in branch B may be affected by loss of *Ret*. Additionally, several established markers of branches A and B (Morarach et al. 2021) are differentially expressed in accordance with expectations (*Etv1, Gal, Nos1,* and *Vip are* higher in cluster 7/branch A; *Bnc2, Dlx5,* and *Ndufa4l2* are higher in cluster 8/branch B).

Both Ret and CFP are downregulated along the branch B trajectory in addition to Gfra1 (Figure 3D), which encodes the canonical RET co-receptor, and Ednrb, which is known to be epistatic with Ret (McCallion et al. 2003; Chatterjee and Chakravarti 2019; Carrasquillo et al. 2002). Interestingly, expression of these same genes is maintained in Ret^CFP/+^ cells along the branch A trajectory, and each exhibits a biased expression for cells along this trajectory (Figure 3D). The exclusion of Ret^CFP/CFP^ cells, and the branch-A-specific persistence of Ret and its canonical co-receptor indicates that commitment to the branch A developmental trajectory, and subsequently, establishment of the enteric neuronal subtypes derived from cells adopting this trajectory, requires continued Ret expression. The downregulation of both Ret and CFP expression along the branch B trajectory, which contains numerous cells of both genotypes (Figure 3A&D), suggests that this trajectory may not require persistent Ret expression. While prior immunohistochemical studies found Ret protein expression is down-regulated in emerging glia and maintained in emerging neurons (Young et al., 1999), our scRNA-Seq data indicate that Ret exhibits differential expression between the two branches of differentiating neurons with Ret expression maintained in branch A and down-regulated in branch B, consistent with other recent single cell studies (Elmentaite et al., 2021; Morarach et al., 2021). In aggregate, these results suggest that in the Ret^CFP/CFP^ developing ENS, since branch A is an inaccessible developmental trajectory, and a greater portion of neuronally committed ENCDCs may engage the branch B trajectory.

To validate Branch A inaccessibility in *Ret^CFP/CFP^* ENCDCs, we selected *Cartpt* (branch A/cluster 7) and *Gfra2* (branch B/cluster 8) from the list of differentially expressed genes as markers for *in vivo* validations due to their high expression and high specificity in each branch (Figure 3D). We performed whole-mount immunohistochemistry (IHC) on isolated gut (esophagus, stomach, and intestine) of E12.5 and E14.5, *Ret^CFP/+^* and *Ret^CFP/CFP^* embryos to identify and localize cells from each branch. We probed for CART, the product of *Cartpt*, and CFP to localize branch A cells. In the *Ret^CFP/+^* samples at E12.5 we observe co-expression of CART and CFP in the stomach (Figure 3E) but not in the esophagus (data not shown). By E14.5, we observe co-expression of CART and CFP extending all the way to the hindgut of *Ret^CFP/+^* samples (Figure 3F). Consistent with our scRNA-seq data, we do not observe any CART expression in the stomach of *Ret^CFP/CFP^* samples at either age (Figure 3E-F) nor in the esophagus (data not shown). We further validated the lack of branch A cells within the intestine of *Ret^CFP/CFP^* cells using single-molecule fluorescent in situ hybridization (RNAscope) to assess mRNA expression of the branch A marker genes *Cartpt* and *Gal* (Figure S6B-C). These results demonstrate that cells associated with the branch A developmental trajectory are not produced within the GI tract in the absence of *Ret*.

To determine the spatial distribution of branch B cells we exploited the specific expression of *Gfra2*, which encodes a non-canonical RET co-receptor, in branch B neurons, which is correlated with a decrease in expression of *Gfra1* (Figure 3D). When we probed for GFRA2 and CFP via IHC we did not observe GFRA2 expression in the E12.5 stomach of *Ret^CFP/+^* samples (Figure 3G). We did, however, observe co-expression of GFRA2 and CFP restricted to cells along the extrinsically innervating nerve fibers of the stomach of *Ret^CFP/CFP^* mice at this age (Figure 3G), confirming that branch B neuronal development trajectory is indeed accessible in the absence of *Ret*. At E12.5 we did not observe GFRA2 outside the extrinsically innervating nerve fibers of the *Ret^CFP/CFP^* stomach, nor in the intestines of either genotype (data not shown), consistent with the previously described migratory defect and failure to effectively colonize the ENS in the *Ret^CFP/CFP^* mice. We were able to detect CFP^+^/GFRA2^+^ cells (putative branch B neurons) in the esophagus of both genotypes, as well as CFP^+^/GFRA2^-^ (non-neuronal ENCDCs) and CFP^-^/GFRA2^+^ (maturing branch B neurons) cells (Figure S6D), indicating that branch B cells are spatially restricted to the esophagus in E12.5 *Ret^CFP/+^* samples and to the esophagus and extrinsic nerve fibers along the stomach in E12.5 *Ret^CFP/CFP^*. By E14.5, CFP^+^/GFRA2^+^ branch B maturing neurons are observed throughout the intestine in the *Ret^CFP/+^*; however, CFP^+^/GFRA2^+^ cells remain restricted to the extrinsically innervating nerve fibers along the stomach in the *Ret^CFP/CFP^* mice (Figure 3H). Consistent with this difference, at E12.5 we see proportionally more *Ret^CFP/CFP^* cells in cluster 8 (branch B) than *Ret^CFP/+^* within our scRNA-Seq dataset (9/37 *Ret^CFP/+^*, 28/37 *Ret^CFP/CFP^*). By E14.5 however, *Ret^CFP/+^* are the predominant cells in cluster 8 (21/33 *Ret^CFP/+^*, 12/33 *Ret^CFP/CFP^*). This could be related to an overall reduced number of E14.5 *Ret^CFP/CFP^* cells in the intestine, or reduced capture efficiency as fluorescence decreases over biological time along this trajectory due to the downregulation of *Ret (*and by proxy *CFP*) expression as cells progress along branch B (Figure 3D).

These data indicate that branch A and branch B represent two transcriptionally distinct developmental trajectories of maturing ENS neuronal subtypes that colonize the length of the intestines in the *Ret^CFP/+^* mice. Branch A neurons are never produced in the *Ret^CFP/CFP^* mice. Furthermore, Branch B neurons are produced, at a potentially faster developmental rate, but remain restricted to the extrinsically innervating nerve fibers throughout development in the *Ret^CFP/CFP^* mouse. As *Ret* is required for cells to migrate into the developing ENS (Figure 2D) (Hirst et al. 2018) it follows that branch A, which appears to require *Ret*, would be inaccessible while branch B, which is *Ret*-independent, remains accessible in the absence of *Ret*.

### HSCR-associated genes are expressed across different cellular states in the developing ENS

While coding and enhancer LoF variants in *RET* contribute to ∼50% of cases of Hirschsprung disease, HSCR is also associated with rare coding pathogenic alleles (PAs) in 23 additional genes (*GDNF, GFRA1, NRTN, SOX10, EDNRB, EDN3, ECE1, ZEB2, PHOX2B, KIFBP, TCF4, L1CAM, IKBKAP, SEMA3C, SEMA3D, NRG1, ACSS2, FAM213A, ADAMTS17, ENO3, SH3PXD2A, SLC27A4* and *UBR4*) (Tilghman et al. 2019; Chatterjee et al. 2016a; Kapoor et al. 2015; Chatterjee et al. 2021). Many of these genes are a part of the *RET-EDNRB* gene regulatory network (GRN) (Chatterjee et al. 2016a). To assess the distribution of HSCR-associated genes, we investigated the expression pattern of all 24 genes in our data to determine their co-expression patterns and their specific effects due to *Ret* LoF (Figure 4A). Three HSCR-associated genes (*Gdnf, Edn3, and Nrtn*) are not expressed in our ENCDC dataset as they are predominantly expressed from the surrounding gut mesenchyme. Of the remaining 21 expressed genes (Figure S7A-C), six are significantly differentially expressed (monocle likelihood ratio test, q<0.1, Figure 4B-C, Table 6A) between *Ret^CFP/+^* and *Ret^CFP/CFP^* cells (*Gfra1, Nrg1, Phox2b, Ednrb, Sema3c*, and *Sox10)*. To identify cell-type-specific differential effects from *Ret* LoF we further tested the combinatorial effect of genotype and cell type. Three HSCR-associated genes (*Gfra1, Sema3d* and *Ednrb*) exhibit significant cell-type-specific differential expression (monocle likelihood ratio test, q<0.05, Figure 4B-C, Table 6D). Both *Gfra1 and Ednrb* reach statistical significance in both tests, suggesting a cell autonomous effect of *Ret* LoF on the expression of these two genes. Alternatively, four HSCR- associated genes (*Nrg1, Phox2b, Sema3c*, and *Sox10*) are globally differentially expressed but are not differentially expressed with respect to genotype:cell type interactions (monocle likelihood ratio test, q<0.05; Table 6A,D).

**Figure 4:**
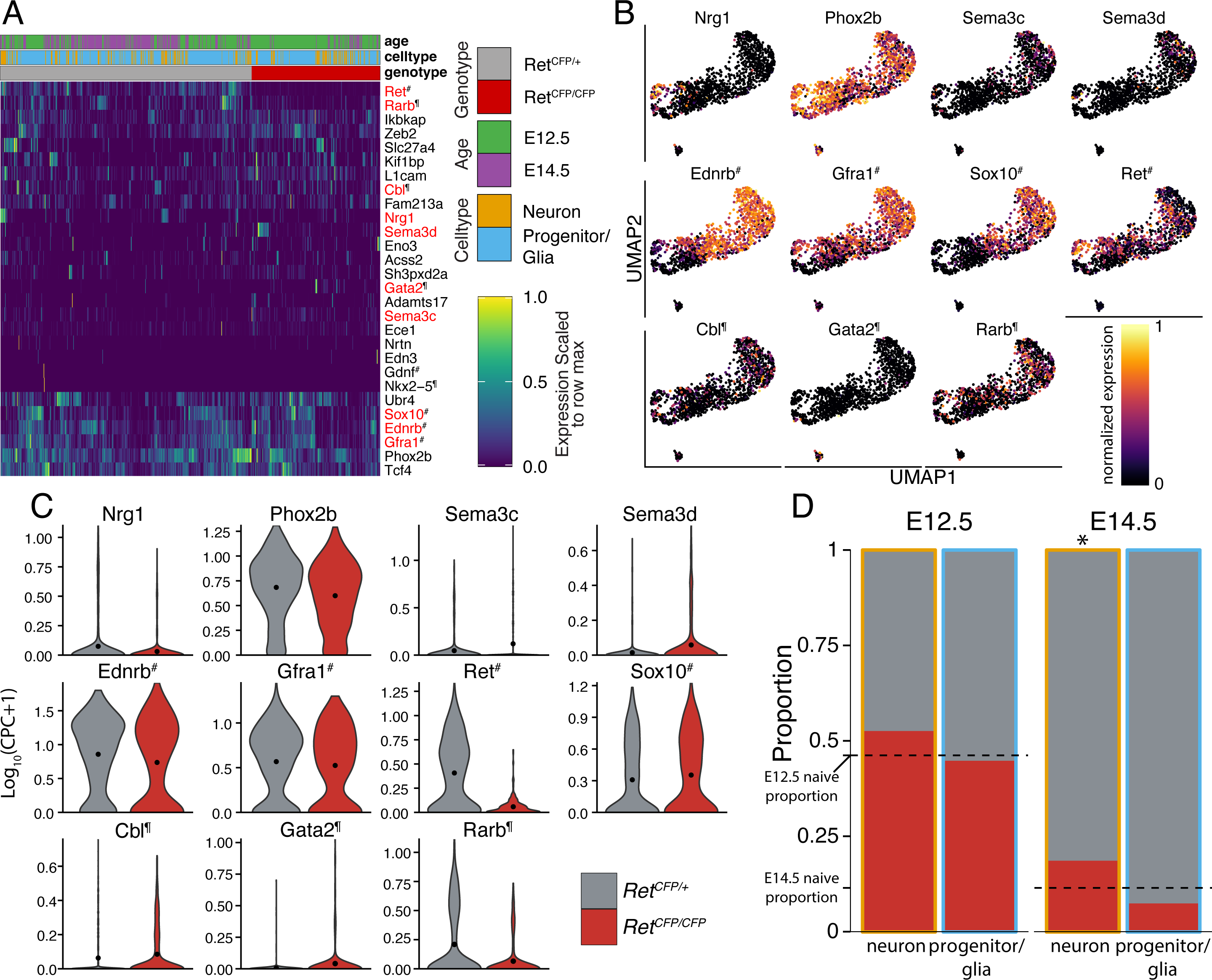
Cellular diversity of Hirschsprung disease (HSCR) genes. (A) Heatmap of expression levels for 24 HSCR-associated genes across all isolated ENCDCs. Significant differentially expressed genes are highlighted in red. (B) UMAP embeddings showing the expression of 11 genes that harbor pathogenic alleles in HSCR patients or are part of the *RET-EDNRB* GRN and are differentially expressed (monocle likelihood ratio test q<0.05) with respect to genotype or with respect to genotype-cell type interactions. Unannotated genes are HSCR-associated but not part of the *RET-EDNRB* GRN, # indicates genes that are both associated with HSCR and are part of the RET-EDNRB GRN, and ¶ indicates genes that are part of the *RET-EDNRB* GRN but have not been associated with HSCR. (C) Violin plots of these 11 genes show expression with respect to genotype, with mean expression value indicated by a dot. (D) Barplot showing the proportion of neurons and progenitors/glia by genotype at E12.5 and E14.5. Dotted lines indicate the naive genotype proportion at E12.5 and E14.5. Asterisk indicates a significant deviance from the naive proportion (Chi-squared test for equal proportions, p<0.05).

We have previously shown that the proportion of cells by genotype deviates significantly from the naïve proportion in seven of the nine clusters in our data (Figure 1D), we also observe that genotype proportions deviate significantly from expectation in E14.5 neurons and approach statistical significance in E14.5 glia (chi-squared test for equal proportions; neurons/p=0.0158, glia/p=0.0702) (Figure 4D). Several of the HSCR-associated genes (*Ednrb, Gfra1, Elp1, Sox10, Zeb2*) are expressed predominantly in progenitor/glia cells (Figure S7A,C). *Phox2b, Tcf4,* and *Ubr4* are the three within this set that are ubiquitously expressed across all clusters in our embedding (Figure S7A,C). *L1cam* is enriched in both maturing neurons and maturing glia, but exhibits reduced expression within the cycling progenitor population of cells (Figure S7A,C). *Ece1* is predominantly expressed in the population of maturing glial cells. These results suggest that some of the observed differential expression of HSCR-associated genes when observed at a global level, but not when assessed within discrete cell states, may be primarily due to differences in cell type proportions across genotypes. Finally, similar to the observed persistence of *Ret* expression along Branch A, a subset of HSCR-associated genes are differentially expressed (monocle likelihood ratio test; q<0.05, Table 6C) between the branches, with higher expression in branch A relative to branch B (*Ednrb, Gfra1, Nrg1*, and *Ret*), suggesting their potential role in the establishment of Branch A identity in conjunction with *Ret*.

Despite significant genetic heterogeneity, ∼67% of HSCR patients have mutations in the genes of the *RET*-*EDNRB* GRN (Tilghman et al. 2019). This GRN includes bidirectional transcriptional feedback between *RET and EDNRB,* and the transcription factors *SOX10, GATA2, RARB,* and *NKX2-5*, as well as *GFRA1* and its physiological ligand, *GDNF*, and the ubiquitin ligase *CBL*, which breaks down phosphorylated RET (Chatterjee and Chakravarti 2019; Chatterjee et al. 2016a, 2019). To ascertain whether this feedback is cell autonomous or not we looked at the expression of these genes in our dataset. *Nkx2-5* and *Gdnf* are not expressed in our ENCDC dataset. Of the remaining six genes, four (*Gata2, Gfra1, Rarb, and Sox10*) are significantly differentially expressed between *Ret^CFP/+^* and *Ret^CFP/CFP^* ENCDCs (Table 6A), with *Gfra1* and *Rarb* more highly expressed in *Ret^CFP/+^* cells and *Sox10 and Gata2* more highly expressed in *Ret^CFP/CFP^* cells, although these moderate differences in expression may also reflect a disproportionate distribution of ENS cell types within our dataset with respect to genotype (Figure 4C-D, Figure S7C). Finally, with the exception of *Sox10*, which rapidly decreases expression during neuronal commitment, and *Gata2*, which is specific to cluster 9, *Ednrb, Rarb, Gfra1*, and *Cbl* all exhibit persistent expression along the branch A neuronal developmental trajectory (Figure 4B, Figure S7C), consistent with a role for the *RET-EDNRB* GRN in establishing branch A identity. The expression of *Sox10* exclusively within the progenitor population explains a previous observation that *Sox10* haploinsufficiency only affects maintenance of progenitor cells and not neurons (Paratore et al. 2002).

These results highlight the diverse cellular states in which HSCR-associated genes are expressed, suggesting that their roles in HSCR disease etiopathology may involve modulation of different aspects of ENCDC proliferation and development. Collectively however, expression of *RET-EDNRB* GRN genes is affected by the loss of Ret signalling and is persistent along the Branch A trajectory, which is inaccessible in the *Ret^CFP/CFP^* mouse.

### Ret LoF reduces the number of actively proliferating ENCDCs in the developing gut

Reduction in proliferation capacity and precocious differentiation as a result of LoF of other genes within the developing ENS have been proposed as significant drivers of colonic aganglionosis (Fujiwara et al. 2018; Bergeron et al. 2016). Given the previously established role for *RET* as a proto-oncogene (Grieco et al. 1990; Mulligan 2014) and its association with increased proliferation in other context (Park and Bolton 2017; Perea et al. 2017; Gattelli et al. 2013), we asked whether *Ret* LoF was associated with significant differences in the proliferation rate of ENCDC progenitors. Differential gene expression analysis with respect to both genotype and cell type identified 331 genes (monocle likelihood ratio test, q<0.05; Figure S8A, Table 6D), consisting of several genes associated with the mitotic cell cycle (C*dkn1c, Cdk1, Top2a, Ccnd1, Cenpk, Cenpe, Cenpf, Ccna2, Ccnb1, Ccnb2*), DNA replication (*Pcna, Mcm3, Mcm5, Mcm6, Mcm10, Gmnn*), and cell cycle regulation (*Rgcc, Mad2l1, Chek2, Rad51ap1*). Gene set enrichment analysis of this differential gene list identified several cell cycle-associated gene sets including mitotic cell cycle processes, cell division, DNA replication, regulation of cell cycle as top scoring Gene Ontology Biological Processes (GO:BP) (hypergeometric test, q<0.05; Table 7D). To focus on cells that are potentially engaging the cell cycle, we next performed a differential gene expression analysis on just the subset of ENCDCs identified as progenitors or glia within the contiguous manifold (i.e. excluding ENCDCs committed to a postmitotic neuronal fate and excluding cluster 9). We identified 364 genes with significant differential expression with respect to genotype (monocle likelihood ratio test, q<0.05; Figure 5A, Table 6E), and again this list was enriched for gene sets associated with cell cycle regulation, in addition to gliogenesis, cell fate commitment, and neural crest cell differentiation (hypergeometric test, q<0.05; Figure 5B, Table 7E). Thus, both cell cycle occupancy and differentiation state, two processes that are intrinsically linked during developmental fate specification, are disrupted in the *Ret^CFP/CFP^* progenitor population relative to *Ret^CFP/+^*.

**Figure 5:**
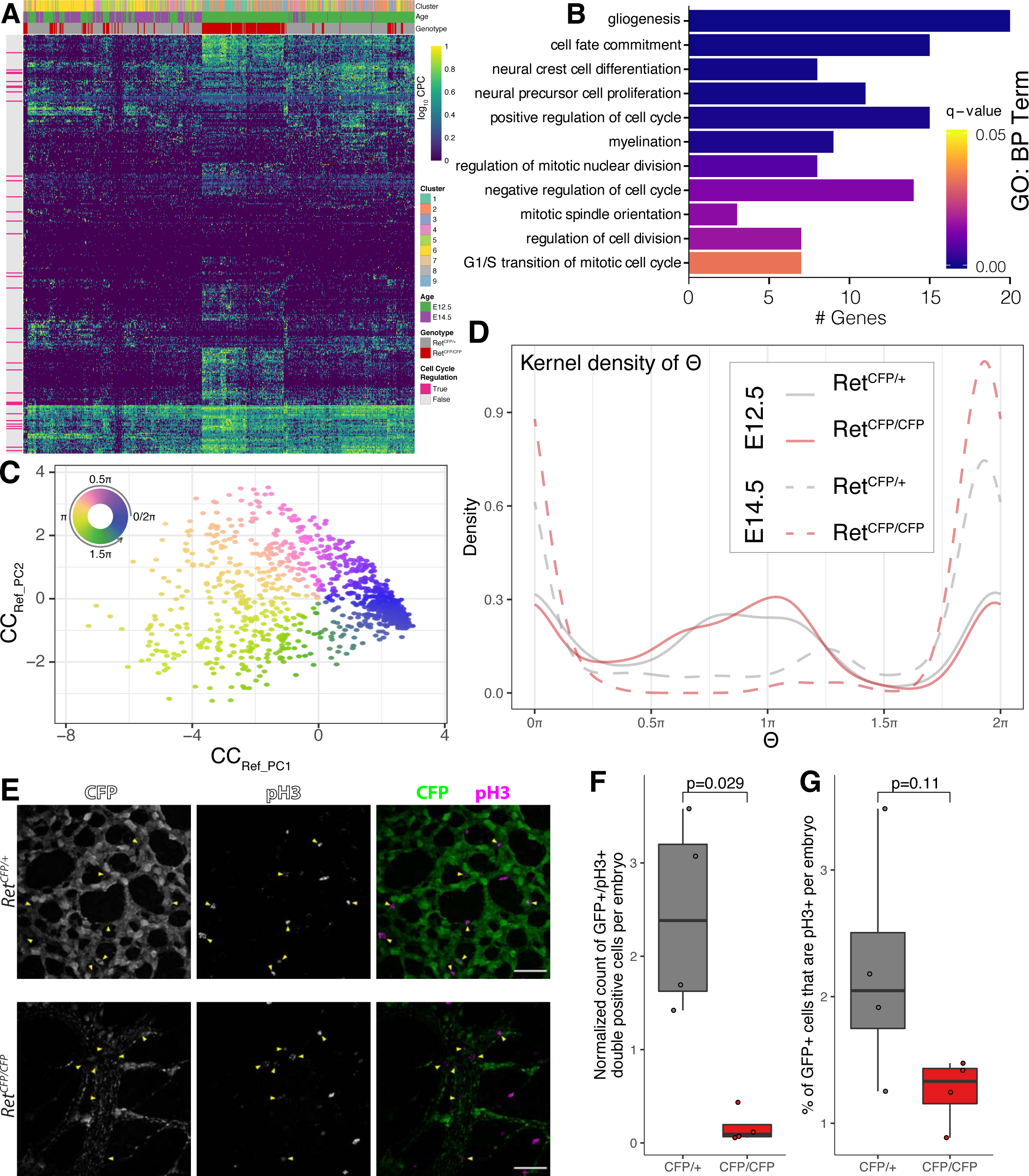
*Ret* LoF results in reduced cell cycle occupancy and significant reduction in the number of actively proliferating ENCDC progenitors. (A) Heatmap of significant differentially expressed genes between *Ret^CFP/+^* and *Ret^CFP/CFP^* within the non-neuronal subset of isolated ENCDCs. (B) Barplots of significant biological process gene ontology terms from gene ontology enrichment analysis of genes with significant differential expression between genotypes in non-neuronal ENCDCs. (C) Tricycle embedding of ENCDCs to estimate cell cycle position. (D) Kernel density estimates for cell cycle position (!) for each genotype at each age. Proliferative phase of the cell cycle (non-G0/G1) is between 0.25” and 1.75”. (E) Representative immunofluorescence staining of the proliferation marker pH3 and CFP in the stomach of E12.5 *Ret^CFP/+^* and *Ret^CFP/CFP^* mice. Yellow arrowheads indicate CFP^+^/pH3^+^ cells. 40X magnification, scale bars are 20 μm. (F) Box & whiskers plot showing the quantification of CFP^+^/pH3^+^ double positive cells per embryo, normalized by the total fields of view analyzed per embryo. (G) Box & whiskers plot showing the quantification of the % of CFP^+^ cells with pH3^+^ staining in the GI tract of E14.5 *Ret^CFP/+^* or *Ret^CFP/CFP^* mice.

We next asked whether we could identify any significant differences in cell cycle state assignments for ENCDCs in the absence of Ret. We used the tricycle package (Zheng et al. 2021) to project the 1,003 single ENCDCs into the default reference cell cycle space provided (Figure 5C). This analysis allows us to estimate the continuous position of each cell along the cell cycle (Θ) (Table 11), comparisons of which by genotype and age (Figure 5D) suggests that at E12.5, there are minimal differences in cell cycle occupancy between genotypes (Kolmogorov-Smirnov test; p-value = 0.660), but by E14.5 there are consistently fewer *Ret^CFP/CFP^* cells engaging the cell cycle than *Ret^CFP/+^*(Kolmogorov- Smirnov test; p-value = 8.28x10^-3^).

To further assess whether *Ret^CFP/CFP^* ENCDCs exhibit differential cell cycle occupancy relative to *Ret^CFP/+^* we next sought to infer cell cycle stage annotations for cells in our dataset using a projection-based approach with an independent, biologically relevant data source. Using the scCoGAPS patterns previously learned from publicly available data, and on which we trained the random forest classification model, we observed high usage of three patterns within a population of cycling glia. Gene ontology analysis of pattern markers revealed these patterns correspond to different proliferative phases of the cell cycle (hypergeometric test, q<0.05, Table 4A&B). Pattern 40 was associated with DNA replication, corresponding to S phase; pattern 41 associated with cilium assembly, organization, and segregation, and attachment of spindle microtubules to kinetochores, corresponding to prophase or metaphase (early M phase); and pattern 42 associated with chromatid segregation and nuclear division, corresponding to anaphase or telophase (late M phase). To predict the cell cycle stage for individual cells in our ENCDC data, we scaled projected pattern weights linearly within each pattern to range between 0 and 1, and used this as a measure of cell cycle stage occupancy probability (Figure S8C&D). We tested all six possible comparisons of the four interaction conditions of genotype and age per pattern for differences in probabilities using two-sample Kolmogorov-Smirnov tests on the empirical cumulative density functions (Figure S8E), and used the Bonferroni method for multiple testing correction. Across the three patterns, E14.5 cells have significantly lower proliferative cell cycle stage occupancy probabilities than E12.5 cells of the same genotype (S phase: *Ret^CFP/+^* Bonferroni adjusted P ≤ 10^-15^, *Ret^CFP/CFP^* /04266516^-15^; early M: *Ret^CFP/+^* /01234516^-15^, *Ret^CFP/CFP^* /03278516^-13^; late M: *Ret^CFP/+^* p01.38x10^-8^, *Ret^CFP/CFP^* p01.94x10^-9^), consistent with the expectation that as progenitor cells differentiate, the proportion of actively cycling cells will decrease over biological time. Again, cell cycle stage occupancy probabilities for ENCDCs did not differ significantly with respect to genotype at E12.5. By E14.5 however, *Ret^CFP/CFP^* cells have significantly lower occupancy scores across all three patterns (S phase: Bonferroni adjusted p09.66x10^- 5^, early M: p01.97x10^-4^; late M: p00.0193) suggesting overall fewer cycling ENCDCs in the *Ret^CFP/CFP^*.

This may be attributed to an increased differentiation rate, and consequently decreased proliferation rate, in *Ret^CFP/CFP^* cells compared to *Ret^CFP/+^* cells between E12.5 and E14.5.

To validate this predicted decrease in cell cycle occupancy in *Ret^CFP/CFP^* ENCDCs, we next used IHC to assess the number of CFP^+^ ENCDC co-expressing the mitotic marker phosphohistone H3 (pH3) within the developing gut at E14.5 (Figure 5E). We confirmed a significant (Student’s t-test, p<1.0x10^-4^) overall reduction in the number of CFP^+^ cells in the *Ret^CFP/CFP^* mouse model at E14.5 (Figure S8G) consistent with the reduced number of CFP^+^ cells observed in our FACS-collected population (Figure 1A) and previously published studies (Uesaka et al. 2008). We also identified a slight difference in the mean number of pH3^+^ cells at E14.5 (Student’s T-test, p<0.05); however, modest enough to suggest that the overall number of mitotic cells within the developing gut remains unaffected (Figure S8H). This is consistent with a cell-autonomous effect of *Ret* on cell cycle as ENCDCs make up a small portion of all proliferating cells in the developing gut and would therefore not contribute significantly to the overall proliferative capacity of the gut as a whole. The overall number of double positive pH3^+^/CFP^+^ cells was significantly reduced in *Ret^CFP/CFP^* animals (Figure 5F & S7I) (Student’s T-test, p=0.029), and while not statistically significant, we observed a difference in the mean percentage of pH3^+^ cells within the subset of CFP^+^ ENCDCs in the *Ret^CFP/CFP^* ENS at E14.5 indicating that a smaller fraction of developing ENCDCs are actively engaging the cell cycle in the absence of *Ret* (Figure 5G & Figure S8J) (Student’s t-test, p=0.11). In aggregate, these results are consistent with the previously established role of *Ret* in enhancing proliferation and indicate that a loss of *Ret* in the developing ENS may contribute to an early reduction in the progenitor pool, possibly through induction of precocious differentiation, leading to a failure to colonize the more distal region of the developing gut.

## Discussion

The enteric nervous system (ENS), initially derived entirely from the caudally migrating vagal and cranially migrating sacral neural crest cells, contains more than 100 million neurons of a variety of functionally distinct subtypes, highlighting the breadth of the functions it performs (Furness 2008; Rao and Gershon 2016). In contrast to the rest of the peripheral nervous system, the ENS is endowed with the ability to integrate multiple neuronal signals to control gastrointestinal (GI) behavior independently from the brain and spinal cord (Furness 2008; Gershon 2010). One of the critical genes that enables this transition of ENCDCs to diverse types of neurons is *RET*, which encodes a cell surface receptor tyrosine kinase. *RET* LoF leads to colonic aganglionosis and is a known factor in the etiology of Hirschsprung disease.

In spite of the critical role *Ret* plays in establishing the normal ENS, there is a limited understanding of the diversity of *Ret*-expressing cells in the developing GI tract and the precise nature of the cell-type- specific effects mediated by *Ret* LoF. By performing scRNA-seq on an enriched population of *Ret*- expressing cells during ENS development we have generated a *Ret-*dependent fate transition map of ENCDCs as they generate diverse classes of neurons and glia in the mouse GI tract to decipher the specific cellular phenotypes mediated by *Ret* LoF.

### Ret LoF affects the establishment of a specific developmental neuronal trajectory in the developing GI tract

Our cellular reconstruction of ENCDC differentiation demonstrates that *Ret-*expressing cells broadly acquire neuronal and glial fates early during normal development of the GI tract and helps explain why *Ret* LoF leads to total aganglionosis caused by impaired migration, defects in proliferation, and loss of immature ENCDCs (Schuchardt et al. 1994). Our data highlight that *Ret* is required for the induction of an entire neuronal specification trajectory within the developing ENS which has previously been associated with the specific development of at least two distinct subtypes of inhibitory motor neurons: an interneuron subtype, and a population of intrinsic primary afferent neurons (IPANs) (Morarach et al. 2021). Though previous studies have posited that branch A corresponds to a predominantly inhibitory motor neuron (MN) trajectory and branch B corresponds to a predominantly excitatory MN trajectory (Morarach et al. 2021), our data are derived from embryonic timepoints which are too early to directly support this functional distinction between the branches. At E12.5, we observe putative branch B cells (GFRA2^+^/CFP^+^) in the esophagus of the *Ret^CFP/+^*, and in the esophagus and stomach of the *Ret*^CFP/CFP^ mice (Figure 3F, Figure S6C). We also observe putative branch A cells (CART^+^/CFP^+^) in the stomach of *Ret^CFP/+^* mice but not the esophagus, and we do not observe putative branch A cells anywhere in the *Ret^CFP/CFP^* GI tract at E12.5. By E14.5 we observe putative branch B cells in the stomach of the *Ret^CFP/CFP^*, and throughout the gut up to and including the wave front of migration in *Ret^CFP/+^* mice. The earlier establishment of branch B cells within the stomach of the *Ret^CFP/CFP^* is consistent with a model of precocious differentiation we identify in this study. The specific failure to produce cells along branch A in the *Ret^CFP/CFP^* suggests that the subset of mature ENS neuronal subtypes derived from this developmental trajectory may be the most acutely affected in disorders such as HSCR and should be the focus of future studies of other *RET*-associated pathologies in the gut.

We provide evidence that aganglionosis observed in HSCR may arise from a combination of early exit from the cell cycle for ENS progenitors, leading to precocious differentiation, and a failure to produce the full repertoire of mature ENS cell types. We do not detect any transcriptional signatures of cell death, suggesting that at this early time point apoptosis or necrosis may not be a significant contributing factor in *Ret*-deficient aganglionosis. The presence of glial cells in our mouse models, independent of genotype, is consistent with previous observations in HSCR patients that immature glial cells can be detected within the aganglionic portion of the colon (Tani et al. 2017). Our data suggest that *Ret* expression is not required for the establishment of a normal ENS glial identity; however, the observed effects on proliferation and differentiation potential suggest that enteric glia may be a preferred fate in *Ret* LoF models.

### Neuronal fates that use non-canonical RET co-receptors remain accessible in the absence of Ret

A previous scRNA-seq study of the developing ENS at E15.5, when the migration of ENCDC through the length of the gastrointestinal tract is near complete, reports a divergence in maturing neurons that results in two branches of progenitors that produce different cell fates (Morarach et al. 2021). We observe this divergence of the neuronal trajectories as early as E12.5, when ENCDCs are still migrating and undergoing neuronal maturation. While we observe a near complete absence of *Ret*^CFP/CFP^ cells in branch A (cluster 7), it is interesting to note that branch B (cluster 8) contains maturing neurons from both genotypes and is not grossly affected by the loss of *Ret*. As cells progress along branch B there is a general reduction in the expression of *Ret*, *CFP*, and several *RET-EDNRB* GRN genes, including *Gfra1.* Concomitant with this, cells in branch B subsequently upregulate *Gfra2*, which encodes a non-canonical RET co-receptor. Though GFRA1 and GFRA2 are in the same family of receptors, GFRA1 preferentially binds GDNF as its ligand, while GFRA2 preferentially binds the ligand NRTN. It is possible that GFRA2 is binding to residual RET after downregulation of transcription; however, it raises the possibility that GFRA2 complexes with another co-receptor to activate an alternate signaling pathway responding to a different ligand in this *Ret*-independent trajectory. Differential expression of *Gfra* family genes within the aganglionic segment of HSCR patients has previously been observed (Lui et al. 2002), including a specific lack of *GFRA1* expression in neuronal cells within the associated hypertrophied nerve fibers. Additionally, *Ntrn* and *Gfra2* null mice exhibit a reduction in neuron fiber density and neuronal size in the ENS, and in each case, result in the specific loss of cholinergic excitatory myenteric innervation (presumably branch B-derived) within the mouse gut, leaving the nitrergic neuronal population (branch A-derived) largely unaffected (Rossi et al. 1999; Heuckeroth et al. 1999), consistent with the segregation of fates and corresponding gene expression differences described here. In aggregate, these results suggest that the canonical signaling pathway of RET, GDNF and GFRA1 is responsible for the proper establishment of ENS subtypes derived from the branch A developmental trajectory, and that the activity of NRTN-GFRA2 signaling may play a greater role in the establishment of ENS neuron identities along branch B, potentially independent from *Ret*.

### Expression patterns of HSCR-associated genes highlight necessity of disease-relevant cellular context

The specific expression patterns of many genes that harbor pathogenic alleles in HSCR patients within the invading and developing ENCDCs of the ENS should lead us to reassess the nature of aganglionosis in HSCR. Our data highlight the need to better understand the specific cellular states and contexts in which these genes and GRNs operate, as well as the cell autonomous versus cell non- autonomous effects in HSCR. Genes such as *Acss2, Eno2,* and *Adamts17* express very lowly in the ENS population (Figure 4), yet are robustly expressed in both the mouse and human developing GI tract (Tilghman et al. 2019; Chatterjee et al. 2019). Furthermore, while the nature of our study, using only sorted *Ret*-expressing cells is by definition only able to capture cell-autonomous effects, additional non- cell-autonomous effects have been observed in *Ret* LoF studies as well (Bogni et al. 2008). Hence patients carrying pathogenic alleles in the genes expressed in other, non-ENCDC, may contribute to other GI anomalies, and aganglionosis may then be a secondary effect rather than primary. Though verification of this would require detailed patient phenotyping, the shift between neuronal trajectories we have identified suggests developmental mechanisms that may guide future analysis of patient samples.

### Ret LoF leads to proliferation defects in ENS progenitors

During formation of the ENS, the gut itself is increasing dramatically in both size and length, creating a huge proliferative burden on ENCDCs, the progenitors of the ENS. Our in silico estimates of cell cycle stage occupancy in conjunction with an *in vivo* assay of proliferation suggest a decrease in proliferation of ENCDCs in the absence of *Ret*. Even a small reduction in proliferative capacity could produce a significant decrease in population size over the course of the several days it takes to colonize the ENS, such as the absence of CFP+ cells we observe in the *Ret^CFP/CFP^* mouse intestines.

*Ret* deletion in the mouse leads to significant changes in fate acquisition, progenitor proliferation, and ENCDC migration within the developing ENS. It is likely that *RET* deficiency alleles in humans lead to similar phenotypes in distinct subsets of ENS cell types and states in HSCR patients, and that each of these phenotypes may play some combinatorial role in establishing the aganglionosis that is the hallmark of this disease. Furthermore, different *RET*-deficiency alleles and epistatic interactions may affect these phenotypes in different ways, potentially explaining some of the observed variable expressivity and incomplete penetrance of HSCR. Our study highlights the specific roles that *Ret* plays within different cellular contexts of the developing enteric nervous system, and demonstrates that appropriate cellular context is critical for the interpretation of multiple disease-related cellular phenotypes that may emerge from a single causal insult.

## Methods

### Mouse strains

Mice heterozygous for the conditional *Ret* allele (*Ret^fl/+^)(Gould et al. 2008; Uesaka et al. 2008)* were obtained from David Ginty (Harvard University, Boston, MA). Briefly, the floxed *Ret* allele (*Ret^fl^*) was generated by inserting a gene cassette, comprising the floxed human *RET9* cDNA with SV40 intron polyA sequence followed by a cyan fluorescent protein (CFP) cDNA with polyA sequence, into exon 1 of the mouse *Ret* gene. *Ret CFP* knock-in mice (*Ret^CFP/+^*; MGI:3777556) were generated by crossing *Ret^fl/+^* mice to β-actin-Cre mice to remove the *RET9* cDNA This mouse was backcrossed >20 generation to C57BL/6 to maintain the line and all experiments were conducted in this isogenic strain. All mouse experiments were conducted in accordance with the National Institutes of Health Guidance for the Care and Use of Laboratory Animals. All procedures were approved by the Johns Hopkins Animal Care and Use Committee (Protocol number: MO21M56). All mice embryos used were either E12.5 or E14.5 and both male and female mice were used.

### Mouse breeding and genotyping

Homozygous *Ret* null mice are perinatal lethal (P0), therefore we intercrossed *Ret^CFP/+^* mice to generate all possible genotypes. Embryos were genotyped from yolk sac genomic DNA to allow for genotype- and sex-specific analysis. Routine mouse genotyping was performed by PCR with primer pairs and amplicons as follows: wild-type *Ret* allele forward primer 5’- CAGCGCAGGTCTCTCATCAGTACCGCA-3’ and reverse primer 5’- CAGCTAGCCGCAGCGACCCGGTTC-3’ resulting in a 449 bp PCR product in *Ret^+/+^* and *Ret^CFP/+^* embryos (modified from (Jain and Varadarajan 2014)), CFP knock-in allele forward primer 5’ ATGGTGAGCAAGGGCGAGGAGCTGTT-3’ and reverse primer 5’- CTGGGTGCTCAGGTAGTGGTTGTC-3’ resulting in a 615 bp PCR product from *Ret^CFP/+^* and *Ret^CFP/CFP^* embryos. Embryo sex was determined by PCR using yolk sac genomic DNA with the forward primer 5’- CTGAAGCTTTTGGCTTTGAG-3’ and reverse primer 5’-CCGCTGCCAAATTCTTTGG-3,’ mapping to exons 9 and 10 of the *Kdm5c/d* genes resulting in two 331 bp X chromosome-specific amplicons in females and an additional 301 bp Y chromosome-specific amplicon in males (Clapcote and Roder 2005).

### Dissection, Dissociation, Flow Cytometry and Sorting

Whole gut tissue (stomach, fore and hind gut) from *Ret^CFP/+^* and *Ret^CFP/CFP^* male and female embryos at E12.5 and E14.5 were dissociated into single-cell suspensions using Accumax (Sigma, USA). The cells were filtered serially through a 100 μm and 40 μm cell strainer, and centrifuged at 2000 rpm for 5 m. The cell pellet was resuspended in 5% FBS, 4 mM EDTA in Leibovitz L-15 medium for cell sorting using MoFlo XDP (BD Biosciences) with a blue-violet excitation laser for CFP. Single cells were sorted directly into 96 well plates containing 1.9 μL Triton X-100 (Sigma-Aldrich, 0.2% v/v, #T9284), 1 μL dNTP mix (Amresco, 10 mM each, #N557), 1 μL 5’ biotinylated oligo-dT30VN primer (IDT, 10 μM), and 0.1 μL RNasin Plus (Promega, 40 U/μL, #N2615) per well. As a no-template negative control, one well per plate did not receive a cell. Immediately after collection, sorted plates were centrifuged (2500 rpm for ∼5 s), frozen on dry ice, and stored at -80°C until library preparation.

### Single-cell RNA-Seq library Preparation

Libraries were prepared from 2 embryos for each of the 8 conditions (*Ret^CFP/+^* and *Ret^CFP/CFP^*, female and male, E12.5 and E14.5) with two exceptions: libraries were prepared from 1 *Ret^CFP/CFP^* male E14.5 embryo and 4 *Ret^CFP/CFP^* female E14.5 embryos owing to differential viability. Single-cell RNA-seq libraries were prepared following a modified Smart-seq2 protocol (Picelli et al. 2014; Chevée et al. 2018). Reverse transcription and preamplification (20 cycles for E12.5 cells, 22 cycles for E14.5 cells) followed the original Smart-seq2 protocol, with the exception of using 5’ biotinylated NGG template-switching oligonucleotides. Sequencing libraries were generated following the Nextera XT protocol (Illumina, #FC- 131-1096) on a quartered scale, using approximately 200 pg input DNA per sample and 14 enrichment PCR cycles. Nextera XT v2 Index Sets (Illumina, #FC-131-2004) were diluted 1:5 in water prior to use. Libraries were sequenced (paired end, 50 bp read length) on an Illumina HiSeq 2500 to an average depth of 9.91x10^5^ (standard deviation 7.43x10^5^) paired-end reads per cell.

### Data preprocessing

Single-cell RNA-seq libraries were aligned with hisat2 version 2.0.1-beta to a mouse mm10 hisat2 index with the inserted CFP cDNA sequence included in the index. Using samtools 1.2, sam files were compressed to bam files then sorted and indexed. Using Cufflinks v2.2.1 indexed bam files were quantified against the Gencode mouse vM10 assembly and normalized across all 1,369 samples (1,351 cells, 18 negative control wells). 51 samples, both in the bottom 5th percentile for total number of reads and for which less than 50% of reads were mapped, were removed. 22 PCA outliers were removed iteratively. The remaining 1,296 samples (1,280 cells and 16 negative control wells) were normalized again, as before. 261 cells for which less than 1,000 genes were detected at non-zero fragments per kilobase of transcript per million mapped reads (FPKM) were removed, and 4 more cells were subsequently removed as PCA outliers. 28 cells were then removed due to a high ratio of mitochondrial to nuclear gene expression, indicating poor library quality. 1,003 cells and 0 negative control wells passed these quality control measures. The age, sex, and genotype of these 1,003 cells is as follows: 183 embryonic day (E)12.5 female *Ret^CFP/+^*, 164 E12.5 male *Ret^CFP/+^*, 173 E12.5 female *Ret^CFP/CFP^*, 125 E12.5 male *Ret^CFP/CFP^*, 161 E14.5 female *Ret^CFP/+^*, 156 E14.5 male *Ret^CFP/+^*, 37 E14.5 female *Ret^CFP/CFP^*, and 4 E14.5 male *Ret^CFP/CFP^*. FPKM values were transformed to copies per cell (CPC) using the Census algorithm (Qiu et al. 2017). Raw and processed data have been deposited in the NCBI Gene Expression Omnibus (GEO) under accession number GSE192676.

### scRNA-Seq analysis

To reduce the computational burden, analyses of gene expression were limited to expressed genes defined as genes with non-zero CPC values in at least 20 cells, and greater than 1x10^-20^ standard deviation. Differential gene expression tests were performed using the Monocle R package (Trapnell et al. 2014). To reduce noise and batch effects in visualizations a set of 419 genes with high dispersion was defined (Chevée et al. 2018). To determine these genes, we normalized expression by read depth across all expressed genes, then subset cells into five sets according to the pool in which each cell was sequenced. Estimated dispersions were calculated for normalized expression of expressed genes in each set. Overdispersed genes were defined as those with greater than 1.4 times the dispersion fit across all five sets. Uniform manifold approximation and projection (UMAP) was performed on these 419 high- dispersion genes using the Umapr R package (McInnes et al. 2018; [CSL STYLE ERROR: reference with no printed form.]). RNAVelocity estimates were obtained using the Velocyto R package(La Manno et al. 2018). Cluster assignment was performed using the mClust package (Scrucca et al. 2016). Specificity scores for each gene were calculated using the cummeRbund package (Goff et al. 2013). Briefly, Jensen- Shannon (JS) distance was calculated between a vector of mean expression level for each cluster to a unit vector of exclusive expression in each cluster, and specificity is defined as one minus the JS distance.

### Pattern discovery with scCoGAPS

The Coordinated Gene Activity in Pattern Sets (CoGAPS) algorithm employs non-negative matrix factorization to determine patterns of gene expression across cells and returns an amplitude (A) matrix of gene weights by pattern and a pattern (P) matrix of cell weights by pattern. The optimal number of patterns to learn was determined by running several iterations and selecting the maximum number of patterns before observing redundancies and noise. Loom files for publicly available scRNA-seq data from the enteric nervous system of adolescent mice were downloaded for analysis (mousebrain.org). 70 patterns were learned on log-transformed unique molecular identifier (UMI) counts from 11,624 libraries prepared through the 10X Genomics platform using single-cell CoGAPS (scCoGAPS)(Stein-O’Brien et al. 2019). We projected our data into these patterns using projectR (Sharma et al. 2019; Stein-O’Brien et al. 2019), which returned projection weights for each pattern in each cell and each gene. For our data, 25 patterns were learned on a log-transformed, normalized expression matrix of CPC limited to expressed genes, defined above. We projected the publicly available data into these patterns learned on our data again using projectR.

### Random Forest Model

While gene expression patterns were learned on the complete public dataset, which consists of 10,519 glia and 1,105 neurons, we trained the random forest model on all neurons and a subset of 1,660 glia to create a 60:40 glia:neurons training dataset to avoid biasing the model to classifying glial cells. The glia were subsampled such that the distribution of glial cells across clusters in the subset matched the distribution across clusters in the whole glial population. The model was trained on 90% of cells to predict cluster assignment from 70 pattern weights with 10-fold cross-validation resulting in an accuracy of 89.8%. To keep model input consistent, and because pattern weights are constrained to [0,1], projected pattern weights were normalized to [0,1] before being used as input to predict cluster assignment. The parent cell type of each cluster was taken as the predicted cell type for our data.

### Immunofluorescence assay on whole mouse gut

Whole gut tissue (stomach, fore and hind gut) at E12.5 and E14.5 was dissected and washed in ice-cold PBS followed by fixation in 4% PFA at 4°C for 1 h. The fixed tissues were permeabilized and blocked in 5% normal donkey serum + 0.3% Triton-X for 1 h at room temperature (RT). The tissues were incubated with appropriate primary antibodies at 4°C overnight, washed with PBS, then incubated in appropriate secondary antibodies at 4°C overnight (see Table 12 for antibodies and dilutions). Tissues were washed in PBS and and incubated with 0.2 μg/mL of DAPI (4′,6-diamidino-2-phenylindole) for 10 m at RT, followed by incubation for 20 m at RT with CUBIC-1 (Susaki et al. 2014) diluted 1:10 with PBS. Samples were washed with PBS and mounted on Fisherbrand Superfrost Plus microscope slides with ProLong Gold Antifade mounting media and imaged using a Nikon Eclipse Ti confocal microscope.

### RNAscope

Whole gut tissue (stomach, fore and hind gut) at E12.5 was dissected and washed in ice-cold PBS followed by fixation in 1% PFA at room temperature (RT) for several hours. Tissue was dehydrated through an ethanol gradient and cleared with xylenes before embedding in paraffin. Tissue blocks were sectioned on a microtome at 10 μm thickness and mounted on Fisherbrand Superfrost Plus microscope slides. Slides were baked for 1 h at 60°C, allowed to cool to RT, then deparaffinized with xylenes. Fluorescence in situ hybridization was performed following the manufacturer’s protocol for the RNAscope Fluorescent Multiplex V2 Assay using the RNAscope Multiplex Fluorescent Reagent Kit v2 (ACDBio #323100). Briefly, deparaffinized sections were incubated in hydrogen peroxide for 10 m at RT and incubated in 1X Target Retrieval for 15 m at RT, then incubated in Protease Plus for 30 m at 40°C. Sections were incubated in a 50:1:1 mixture of RNAscope Probes CFP (ACDBio #529381), Mm-Cartpt- C2 (ACDBio #432001-C2), and Mm-Gal-C3 (ACDBio #400961-C3) for 2 h at 40°C. Sections were incubated with Amp 1, 2, and 3 for 30 m, 30 m, and 15 m at 40°C, respectively. Next, for each of the three channels sections were incubated in the appropriate HRP for 15 m at 40°C, the appropriate fluorophore for 30 m at 40°C, then HRP blocker for 15 m at 40°C. The fluorophores used were 1:750 dilutions of TSA Plus fluorescein (PerkinElmer #NEL741E001KT), cyanine 3 (PerkinElmer #NEL744E001KT), and cyanine 5 (PerkinElmer #NEL745E001KT). Sections were then incubated in 1X Trueblack Lipofuscin Autofluorescence Quencher (VWR #10119-144) for 1 m at RT to reduce red blood cell autofluorescence. Sections were then incubated in RNAscope DAPI for 30 s at RT and mounted with ProLong Gold Antifade mounting media and imaged using a Nikon Eclipse Ti confocal microscope.

### Image acquisition

Tissues were visualized using a Nikon Eclipse Ti confocal microscope and processed using Fiji. Confocal stacks were taken using a 10X air objective (Figure 2D), 20X air objective (Figure S6B-C), 40X oil objective with Type A immersion oil (Figure 5E and proliferation quantification), or 60X oil objective with Type A immersion oil (Figure 3E-G, Figure S6D). For image presentation, stacks were z-projected to single images using the maximum value for each pixel. For Figure S6B, two fields of view were stitched together (Preibisch et al. 2009).

### Proliferation assay and quantification

To measure cell cycle occupancy we stained replicate *Ret^CFP/+^* and *Ret^CFP/CFP^* whole GI tracts from both male and female mice at E14.5 for phosphohistone H3 (Anti-H3S10p; Millipore Sigma #06-570), an immunomarker specific to cells undergoing mitosis (Skaland et al. 2009), and CFP protein (Anti-GFP; Aves labs #GFP-1010) using the immunofluorescence assay described above. Confocal stack images were acquired on the Nikon Eclipse Ti confocal microscope with a 40X oil objective and Type A immersion oil and quantified using a custom CellProfiler (Carpenter et al. 2006; McQuin et al. 2018) workflow. Briefly, primary objects were identified from each input channel (pH3, CFP) using the Otsu Adaptive threshold strategy with a smoothing scale of 1.34888 and a threshold correction factor upper bound limit of 0.99. The adaptive window size was set at 40. The typical diameter size ranged between 30-120 pixels depending on whether nuclei (pH3) or cell bodies were being identified (CFP). Objects were related and filtered using standard parameters and cells were quantified as either CFP^+^, pH3^+^, or CFP^+^/pH3^+^ double positive. The proportion of dividing ENS precursors for each genotype was determined by estimating the mean % of pH3+ cells within the subset of CFP+ cells for each animal. To directly test the two genotypes, we performed Student’s T-Test on the percentage of pH3^+^ CFP^+^ cells between *Ret^CFP/+^* and *Ret^CFP/CFP^* aggregated to each replicate.

## Acknowledgements

We would like to thank Hao Zhang at the Bloomberg School of Public Health, Johns Hopkins University Cell Sorting facility for help with single cell sorting. E.V. is supported in part by the Johns Hopkins Predoctoral Training Grant in Human Genetics (T32-GM007814-39). L.A.G. and E.V. are supported by an award from the National Institute of Aging (R01AG066768). L.A.G. is additionally supported by the National Institute of Aging (R01AG072305), the National Science Foundation (IOS-1665692), and a Johns Hopkins Catalyst award. SC is supported by a NYU Grossman School of Medicine Startup Grant (59-D-37802-30609-NYUPRJ) and AC is supported by a National Institutes of Health (Eunice Kennedy Shriver National Institute of Child Health and Human Development) R01 award HD028088.

## Author Contributions

Elizabeth Vincent: Investigation, Data Analysis, Writing, Original draft preparation, Review and editing, Validation, Visualization; Sumantra Chatterjee: Conceptualization, Investigation, Writing, Original draft preparation, Review and editing, Supervision. Gabrielle Cannon: Investigation; Dallas Auer: Investigation; Holly Ross: Investigation; Aravinda Chakravarti: Conceptualization, Writing, Review and editing, Supervision, Funding acquisition; Loyal A. Goff: Conceptualization, Data Analysis, Writing, Original draft preparation, Review and editing, Supervision, Funding acquisition

## Supplemental Figures

Figure S1: Uniform Manifold Approximation and Projection (UMAP) embedding of 1,003 cells colored by (A) age, (B) sex, and (C) sequencing batch.

Figure S2: Cluster-specific gene expression profiles (A) Scatter plot of mean gene expression per cluster vs. cluster specificity scores for each of the 9 learned clusters of enriched ENCDCs. (B) UMAP embeddings highlighting the gene expression of top specific cluster markers for each cluster. (C) UMAP embedding showing the expression levels for the ENS glial marker *Ngfr* and the neuronal marker *Tubb3*.

Figure S3: Projection analysis of adult ENS cell types Each panel describes the usage of a single scCoGAPS pattern learned from (Zeisel et al. 2018). For each of the 70 patterns, the left panel is a reduced dimensionality visualization of the glial cells and the middle panel is for the neuronal cells from (Zeisel et al. 2018). Cells are colored based on their usage of the learned pattern. The right panel is a UMAP embedding of the ENCDCs isolated in this study and cells are colored by the corresponding projected pattern weight, highlighting the shared use of specific biological features across both datasets.

Figure S4: Cell type annotation via transfer learning (A) Reduced dimensionality visualization of mature ENS cells from (Zeisel et al. 2018) colored by previously annotated cluster designations. (B) UMAP embedding of sorted ENCDCs colored by the random forest model predictions for corresponding clusters from Zeisel 2018. (C) UMAP embedding of sorted ENCDCs colored by top-level ENS cell type designation. (D) Stacked barplot of cell type proportions for each of the nine learned clusters of ENCDCs. (E) UMAP embedding of Zeisel 2018 mature ENS cells by previously annotated cell type designations.

Figure S5: Learned gene expression patterns across all cells 25 single-cell coordinated gene activity in pattern sets (scCoGAPS) patterns learned from sorted *Ret*- expressing ENCDCs. Each UMAP embedding is colored by the usage of a specific pattern of gene co- expression learned through non-negative matrix factorization.

Figure S6: RNAscope single molecule fluorescence in situ hybridization validation of branch A and branch B cells (A) UMAP embedding of ENCDC cells highlighting previously identified (Morarach et al. 2021) marker gene expression for neuron developmental branches (branch A: *Nos1, Vip, Gal, Etv1*; branch B: *Bnc2, Dlx5, Mgat4c, Ndufa4l2*). (B,C) RNAscope single molecule fluorescence *in situ* hybridization (smFISH) validation of *Gal/Cartpt* in cross section of E12.5 stomach in both *Ret^CFP/+^* and *Ret^CFP/CFP^* mice. The yellow box in (B) indicates the portion isolated in (C). 10X magnification, scale bars are 200 μm (B) and 50 μm (C). (D) Whole mount immunofluorescence validation of branch B cells (GFRA2^+^/CFP^+^) in E12.5 esophagus of both *Ret^CFP/+^* and *Ret^CFP/CFP^* mice. 60X magnification, scale bars are 20 μm.

Figure S7: Expression profiles for Hirschsprung-associated and RET-EDNRB gene regulatory network genes (A) UMAP embeddings showing normalized expression of 21 expressed genes associated with Hirschsprung disease (HSCR). (B) Violin plots showing expression of these same 21 genes by genotype. (C) Violin plots showing expression of these same 21 genes by cluster. (D) UMAP embeddings showing normalized expression of 3 genes in the *RET-EDNRB* GRN that have not been associated with Hirschsprung disease. (E) Violin plots showing expression of these same 3 genes by genotype. (C) Violin plots showing expression of these same 3 genes by cluster. Genes in red are significantly differentially expressed (monocle likelihood ratio test q<0.05) either with respect to genotype or with respect to genotype-cell type interactions. # indicates genes that are shared across the HSCR-associated and *RET- EDNRB* GRN sets.

Figure S8: Cell cycle is modulated by Ret LoF (A) Heatmap of significant differentially expressed genes with respect to the combinatorial effect of both genotype and celltype (i.e. genotype:celltype effect). (B) Barplot of significantly enriched gene ontology terms for the list of genes identified as differentially expressed in A. Ontology coloring matches the row color annotations for each gene in A. (C) UMAP embeddings showing the projected pattern weights for the cell cycle stage-specific patterns (40-42) learned from (Zeisel et al. 2018). (D) UMAP embedding colored by the maximum projection value across each projected cell cycle stage-specific pattern. (E) Empirical cumulative density functions for projection values of cell cycle stage-specific patterns for each of the interaction terms between age and genotype. Significant differences in distributions were determined using the Kolmogorov-Smirnov test (* = p<0.05; ** = p<0.01 after multiple testing correction. (F) Empirical cumulative density function (eCDF) curve across all projected (Zeisel et al. 2018) scCoGAPS patterns. One curve for each of the interaction terms between age and genotype are presented. (G-J) Box & whisker plots showing the distribution of quantifications for pH3 and CFP immunofluorescence staining in E14.5 *Ret^CFP/+^* and *Ret^CFP/CFP^* mouse GI tracts. Each point represents a separate sampled field of view from one of four replicate mice from each condition. Points are colored by replicate embryos. G) Count of CFP^+^ cells, H) Count of pH3^+^ cells, I) Count of double positive CFP^+^/pH3^+^ cells, and J) Percent of CFP^+^ cells that are pH3^+^.

## Tables

Table 1: Distribution of cells before and after QC The age, sex, and *Ret* genotype of each embryo, as well as the number of libraries that were prepared, sequenced, and passed all quality control filters.

Table 2: Gene specificity scores by cluster Specificity scores for each gene (row) in each cluster (column). Specificity scores range from [0, 1], with a higher score indicating greater specificity.

Table 3: Gut atlas scCoGAPS A and P weights Sheet “A-Pmatrix” contains the Pattern (P) matrix of pattern weight values for each cell (row) in each pattern (column) across 70 patterns learned on gut scRNAseq atlas data from Zeisel et al, 2018. Sheet “B-Amatrix” contains the Amplitude (A) matrix of pattern weight values for each gene (row) in each pattern (column).

Table 4: Gut atlas pattern markers Sheet “A-patternMarkers” lists genes that are pattern markers for the 70 patterns in Table 3. Sheet “B- GO_terms” lists the corresponding gene ontology:biological process (GO:BP) terms for which the pattern marker lists are enriched. Sheet “C-GO_term_genes” lists the specific genes associated with each enriched GO:BP term. An entry of “not run” on sheet B indicates the pattern marker gene list for that particular pattern was not long enough (at least 10 genes) after converting to Entrez IDs, therefore gene ontology enrichment was not run on that list. An empty column under a pattern name on sheet B indicates no GO:BP terms reached significance (q<.05) for enrichment in pattern markers of that pattern.

Table 5: ENCDC data projected into gut atlas Projected pattern weights (sheet “projection”) and p-values (sheet “pval”) for the 70 patterns (row) learned on the gut atlas into each cell (column) from the Ret mutant dataset.

Table 6: Differentially expressed gene lists Each sheet presents results from a single differential expression test. At the top of each sheet we report which cells were tested (“cells”), the parameter tested (“test”), the full model formula (“full_model”), and the reduced model formula (“reduced_model”). Gene lists are subset to significantly differentially expressed (q < 0.05) genes.

Table 7: Differentially expressed gene GO lists Each sheet presents results from a single gene ontology (GO) enrichment test, including the number of differentially expressed genes associated with each term (“Count”) and the official gene symbol for those genes (“geneID”) . The sheets in Table 7 report the GO enrichment results for the corresponding differentially expressed gene lists in Table 6. GO enrichment lists are subset to significantly enriched (q < 0.05) terms.

Table 8: ENCDC scCoGAPS A and P weights Sheet “A-Pmatrix” contains the Pattern (P) matrix of pattern weight values for each cell (row) in each pattern (column) across 25 patterns learned on the Ret mutant dataset. Sheet “B-Amatrix” contains the Amplitude (A) matrix of pattern weight values for each gene (row) in each pattern (column).

Table 9: ENCDC patterns correlated with metadata Correlation (sheet “A-correlation”) and the p-value for correlation (sheet “B-correlation-pvalue”) of various cell metadata parameters (row) with pattern weights for 25 patterns learned on the Ret mutant data (column).

Table 10: ENCDC data pattern markers A list of marker genes each of the 25 patterns (column) learned on the Ret mutant data.

Table 11: Tricycle theta estimates Theta estimates from tricycle (“tricyclePosition”) as well as relevant metadata for each cell (row).

Table 12: <u>Antibodies and Concentrations Used</u> A list of all antibodies used in this study.

