## Supplemental Figures for "*Ret* loss-of-function decreases neural crest progenitor proliferation and restricts developmental fate potential during enteric nervous system development"

### Supplemental Figure 1

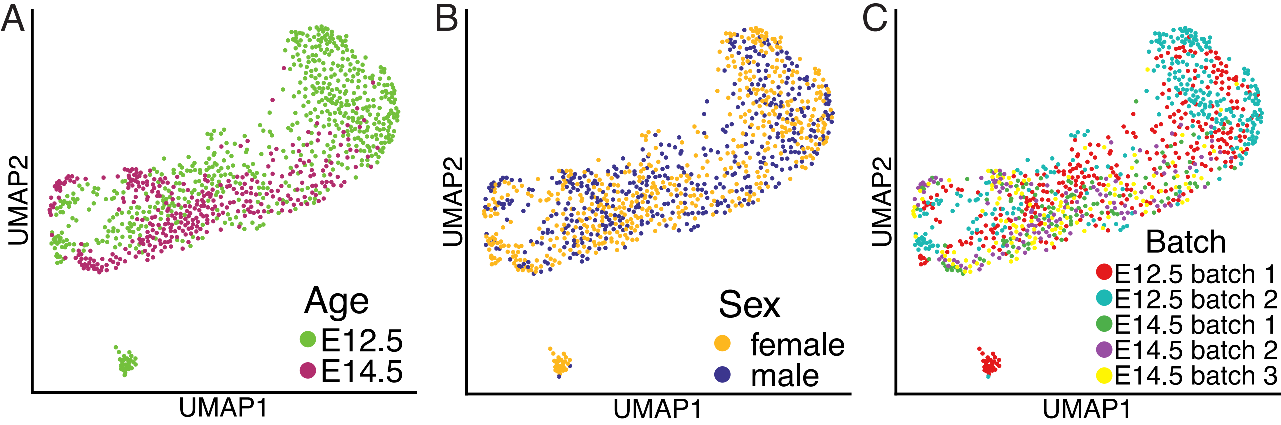

### Supplemental Figure 2

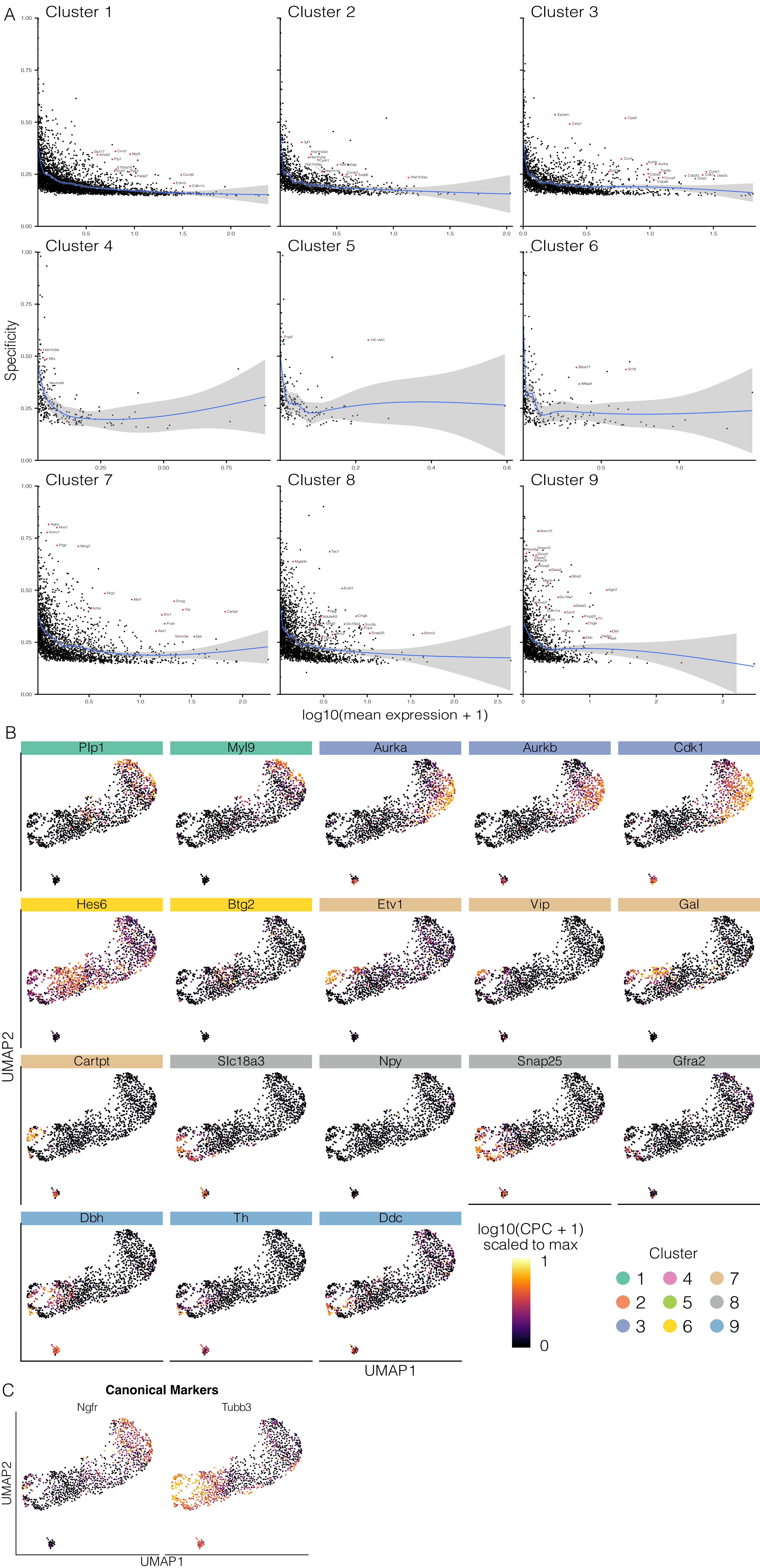

Supplemental Figure 3

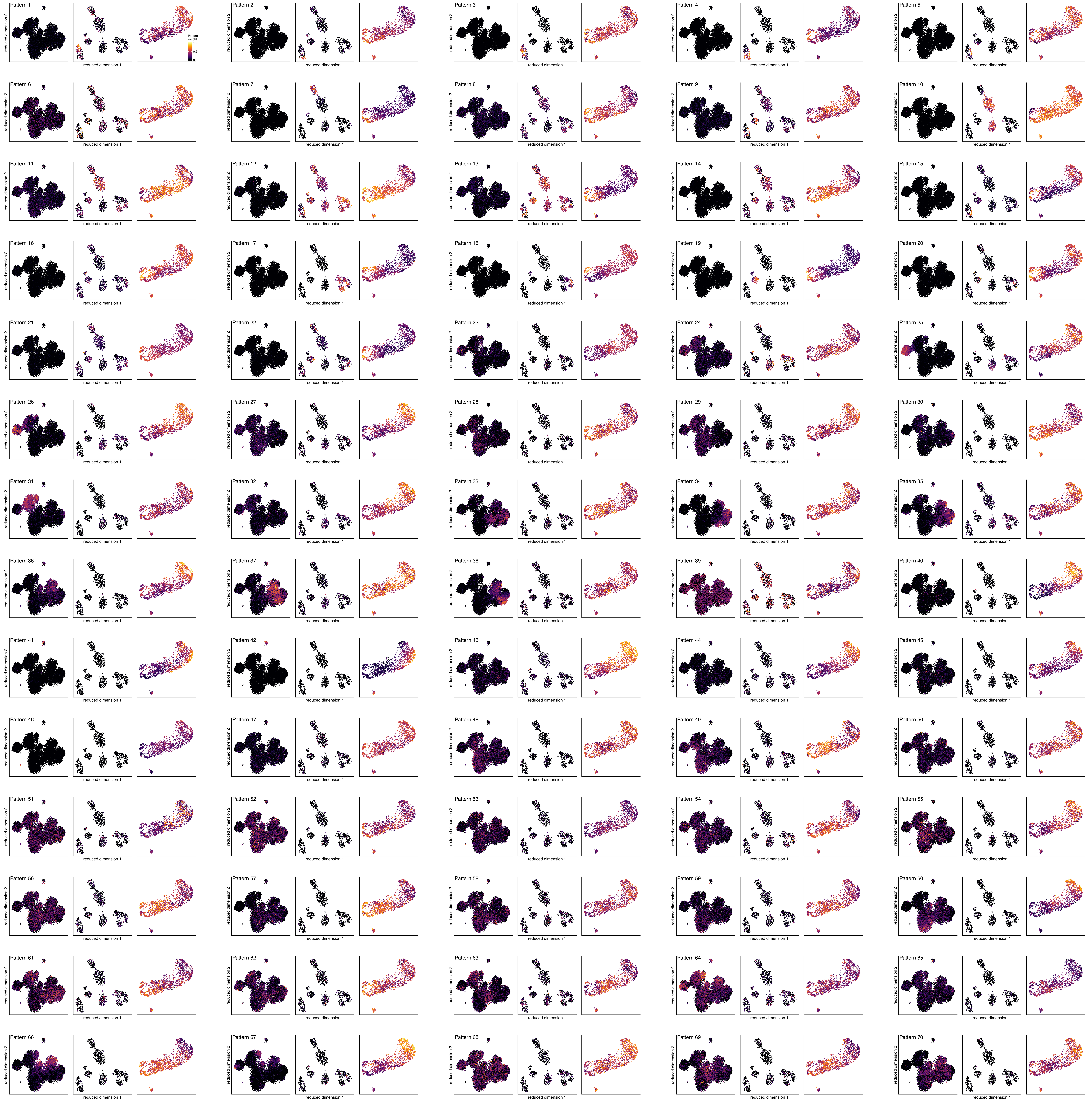

### Supplemental Figure 4

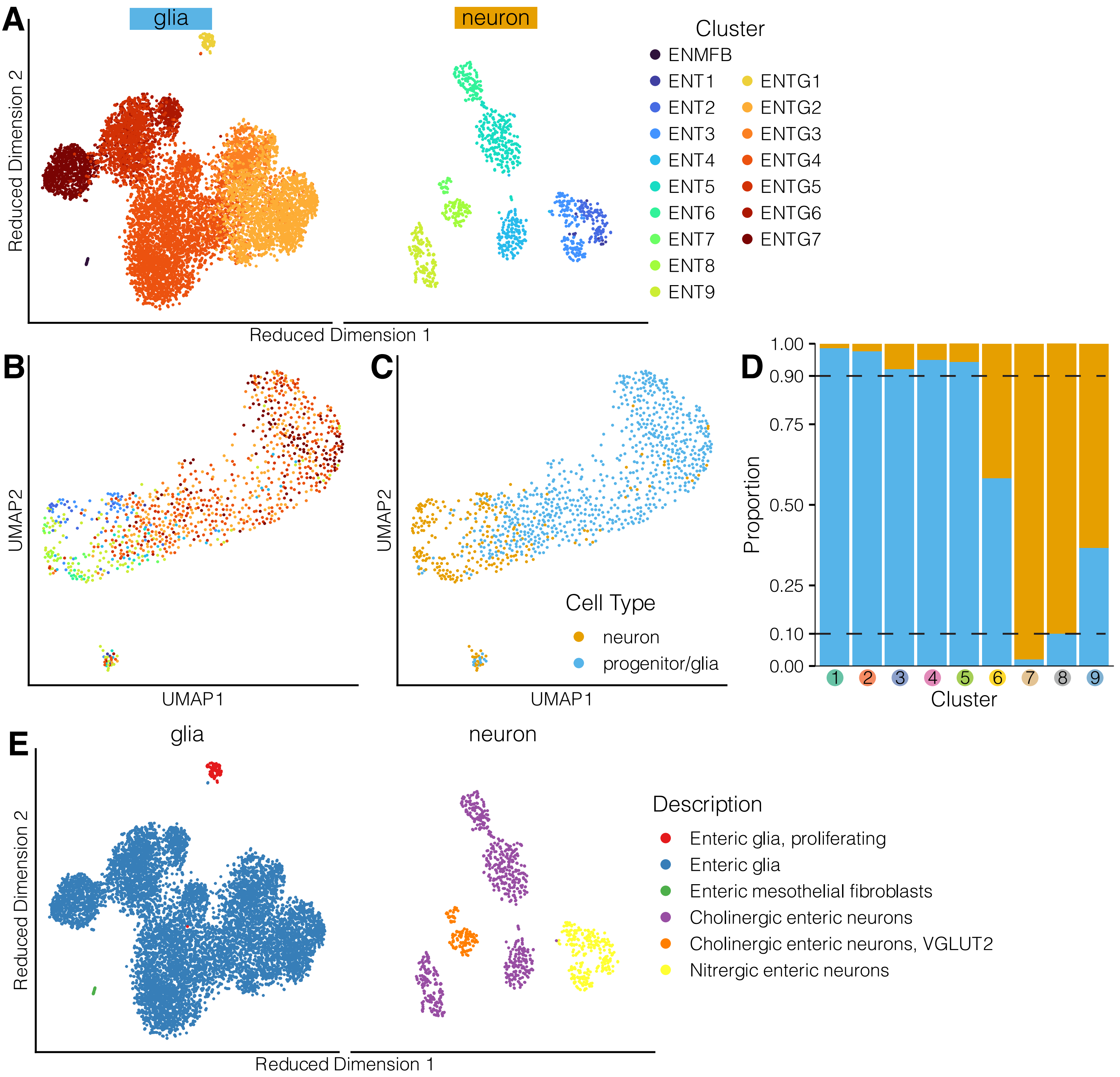

### Supplemental Figure 5

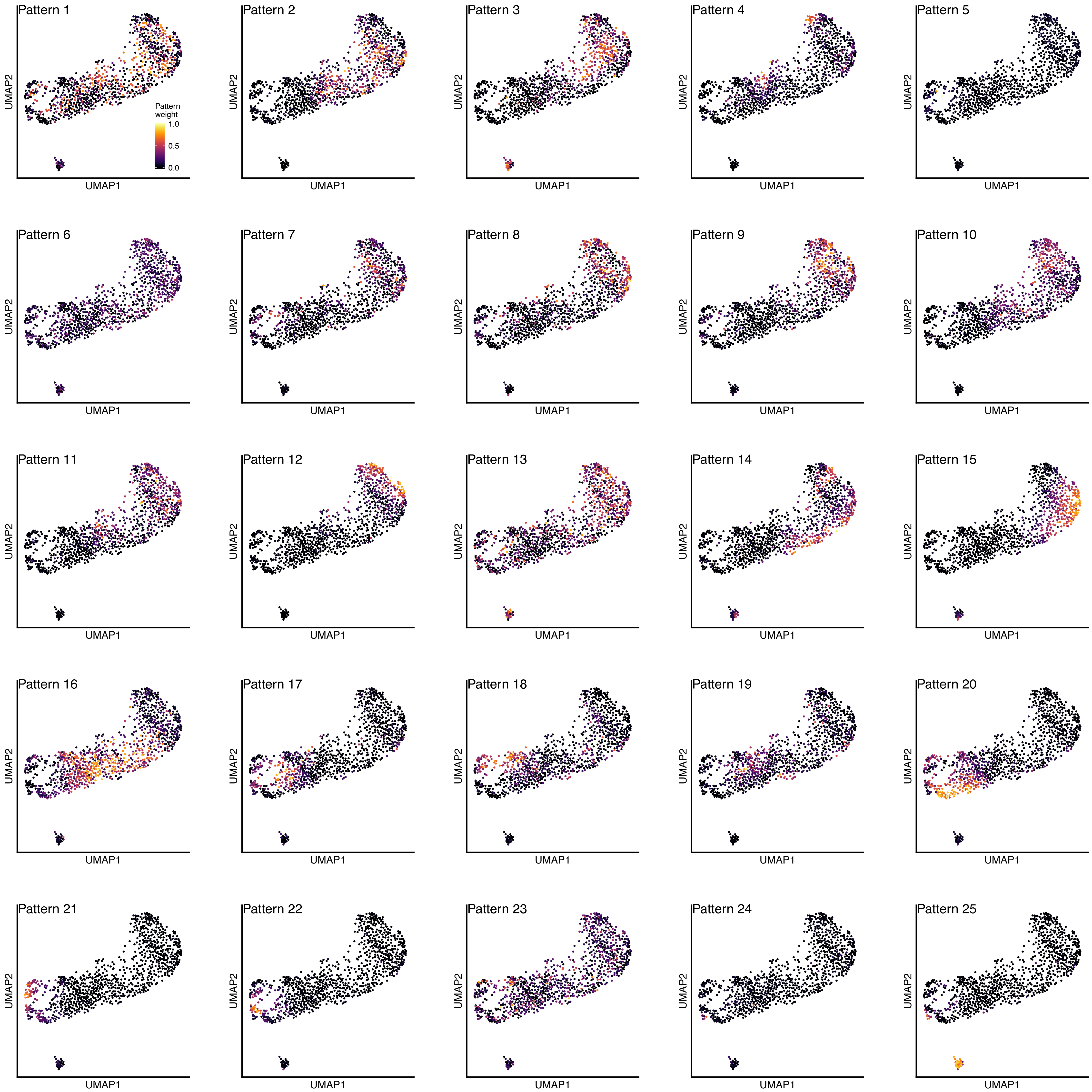

### Supplemental Figure 6

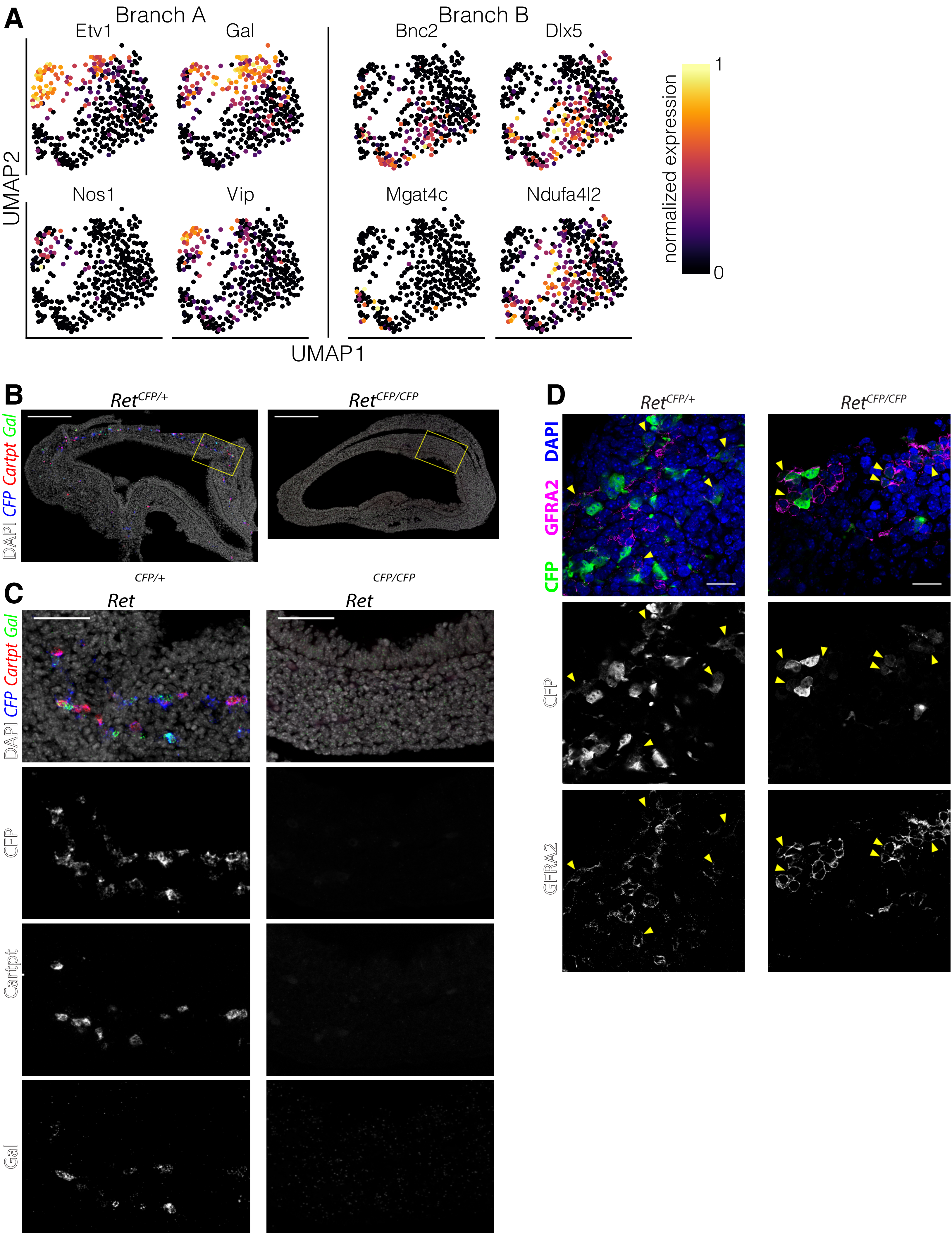

### Supplemental Figure 7

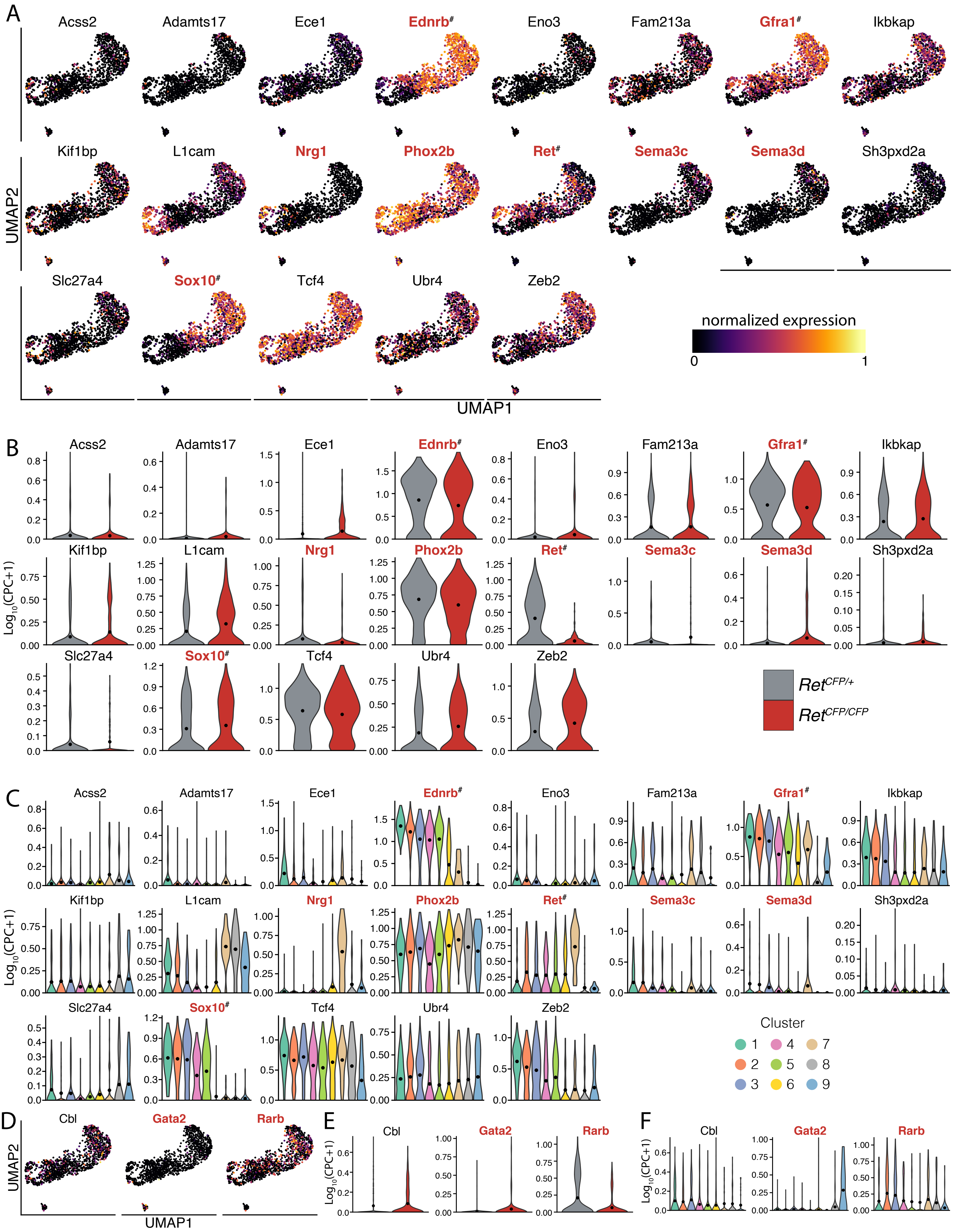

### Supplemental Figure 8

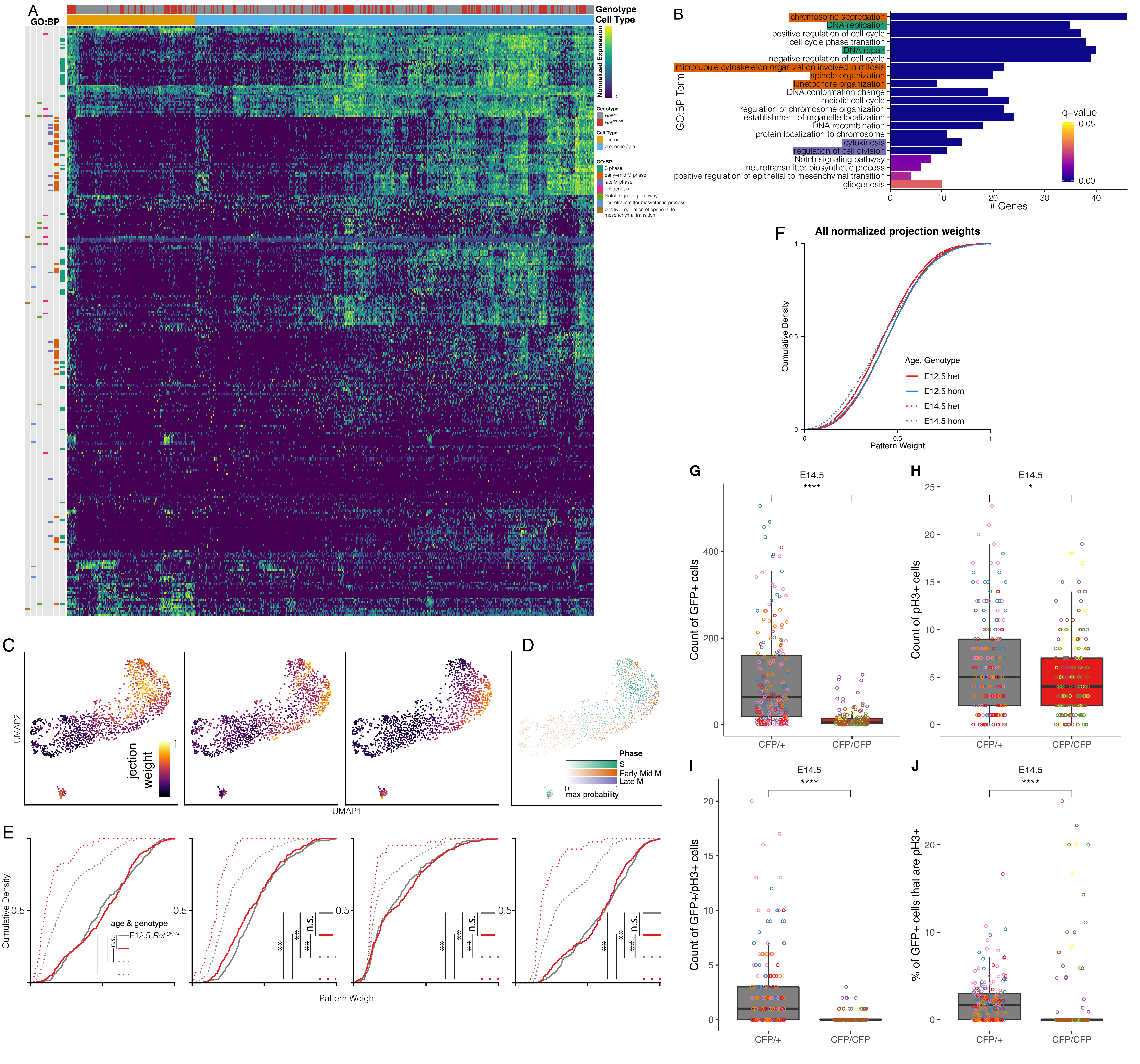
